# Body Fluid Proteomic Landscape of Acute Exercise

**DOI:** 10.1101/2025.05.28.656705

**Authors:** Nigel Kurgan, Jeppe Kjærgaard, Naja Zenius Jespersen, Viriginia Díez-Obrero, Roger Moreno-Justicia, Sarah Elizabeth Heywood, Cecilie Bergmann Lindqvist, Lisa Ottander, Diana Samodova-Sommer, Nina Sloth Nielsen, Cody Garett Durrer, Mathias Ried-Larsen, Simon Rasmussen, Ruth J.F. Loos, Roelof Adriaan Johan Smit, Bente Klarlund Pedersen, Atul Shahaji Deshmukh

**Author notes:** Corresponding authors, Senior and Corresponding Authors Atul S Deshmukh (Lead contact), Novo Nordisk Foundation Center for Basic Metabolic Research, Faculty of Health and Medical Sciences, University of Copenhagen, Blegdamsvej 3B, DK2200, Copenhagen, Denmark, Bente Klarlund Pedersen, Centre for Physical Activity Research, Rigshospitalet, University of Copenhagen, Copenhagen, Denmark, Blegdamsvej 9, DK2100, Copenhagen, Denmark., Nigel Kurgan, Novo Nordisk Foundation Center for Basic Metabolic Research, Faculty of Health and Medical Sciences, University of Copenhagen, Blegdamsvej 3B, DK2200, Copenhagen, Denmark. **Author List Footnotes** Shared first authors (Nigel Kurgan, Jeppe Kjærgaard, Naja Zenius Jespersen).

## Abstract

Physical activity improves health, yet the molecular mechanisms remain partially understood. This study presents a high-resolution, time-resolved atlas profiling 10,127 proteins across plasma, saliva, and urine from healthy adults post-acute exercise. Exercise regulated over 3,000 proteins, revealing distinct, fluid-specific temporal dynamics. By integrating fluid-specific exercise signatures with tissue and disease atlases, we delineated the contribution of tissues and associations to various diseases. Network analysis across body fluids elucidated coordinated remodeling in the extracellular matrix and immune activation orchestrating exercise-induced networks. Many exercise-responsive plasma proteins were robust across age, sex, and exercise modalities, indicating a conserved systemic signature. Integration with genetic data established exercise-regulated proteins as modulators of metabolic traits and identified over 200 targeted by approved drugs, highlighting their impact on disease-relevant pathways. This comprehensive atlas, available as an open-access resource https://cbmr.ku.dk/research/research-groups/deshmukh-group/shiny-apps/, advances our molecular insight into exercise adaptations and enables exerkine discovery, biomarker development, and pharmacological exercise-mimetic strategies.

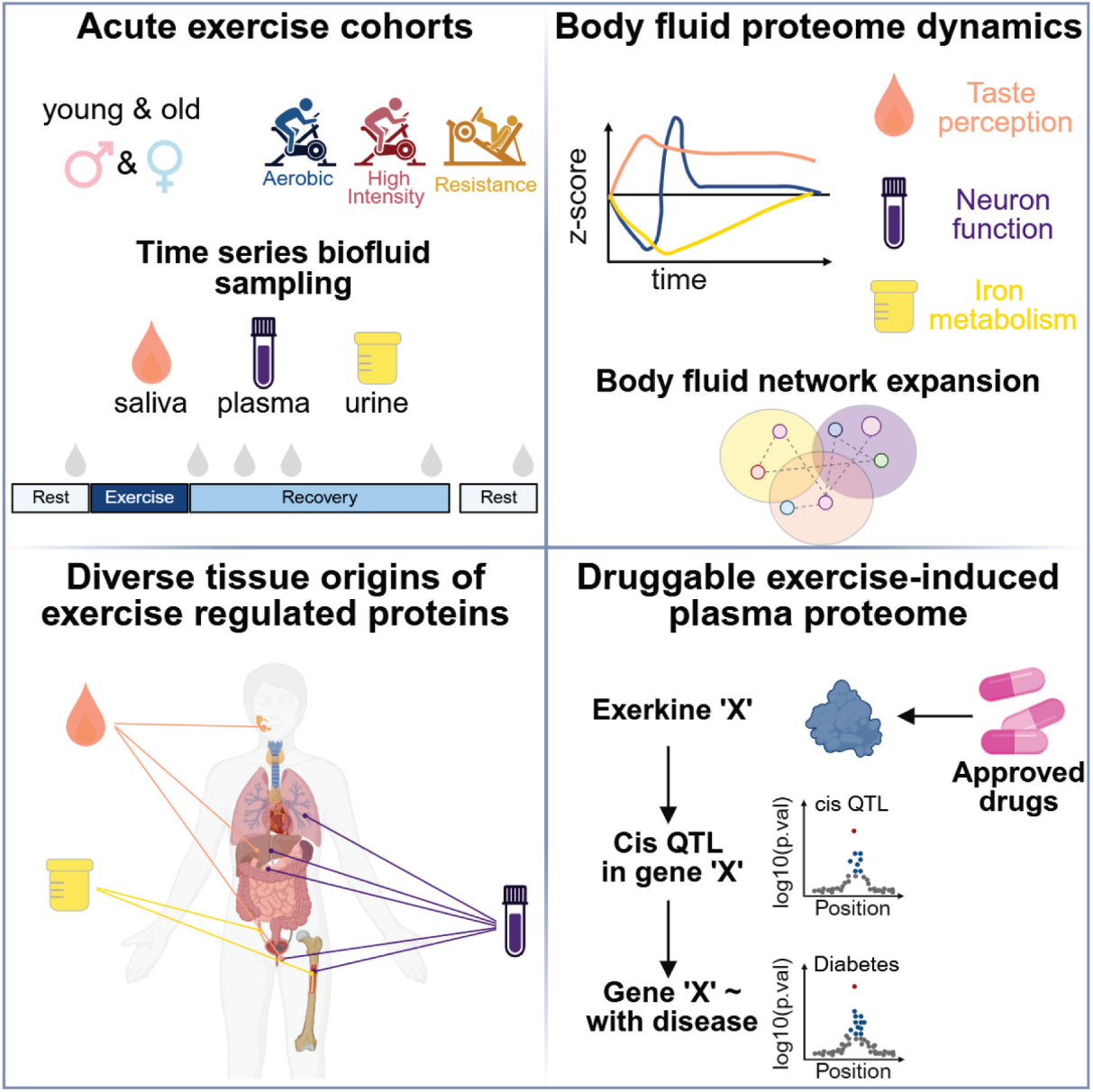

**Highlights:**

- Exercise induces robust and distinct changes across body fluid proteomes
- Tissue remodeling and immune activation drive exercise-induced network expansion
- ∼1,000 exercise-regulated plasma proteins are age, exercise mode, or sex-specific
- Genetic inference identifies druggable exerkines that regulate health and disease

## Introduction

Physical activity plays a vital role in reducing the risk of chronic diseases and promoting longer, healthier lives^1^. However, relatively modern lifestyles have led to a dramatic decline in physical activity levels^2,3^, leaving many people at risk of associated non-communicable diseases, including type 2 diabetes, dementia, cardiovascular disease, and certain cancers^1,4^. Thus, physical inactivity stands as a pressing global health issue^5^, substantially contributing to disease burden and premature mortality^1^. In contrast, 150 min of moderate to vigorous physical activity or exercise per week is associated with a reduced mortality rate of ∼33%^6^, decreased non-communicable disease risk, and increased quality of life through improvements in musculoskeletal function and mental health^7^. Despite this relationship being well established^6^, the molecular mediators responsible for these robust effects of repeated bouts of acute exercise remain unknown.

Molecules secreted during acute exercise, termed ‘exerkines,’ are hypothesized to mediate numerous long-term health benefits of physical activity^8,9^. Exerkines encompass RNAs, proteins, and metabolites/lipids and originate from diverse organs and cell types^10^. Elucidating these mediators may inform the development of targeted exercise interventions and potential therapeutics. Considering that proteins constitute the functional unit of cells and represent the predominant targets for pharmacological interventions, greater emphasis should be directed towards investigating their exercise responsiveness. The most well-characterized example is the myokine IL-6, a cytokine released during acute exercise^11^ that regulates energy allocation in muscle tissue^12^ and mediates long-term adaptations to exercise training^13^. Although the molecular response in blood or serum to acute exercise has been extensively studied^14–18^, these foundational studies were constrained by proteomic coverage, lacked insight into tissue-specific contributions to the exercise-regulated proteome, and did not integrate genetic evidence that could clarify their roles in health and disease. Furthermore, a critical knowledge gap exists regarding exercise-induced alterations in alternative body fluids, such as saliva^19^ or urine^20,21^, which may yield unique or complementary insights into tissue-specific responses to exercise.

Assessment of the urinary proteome can reflect renal clearance of plasma-derived exerkines, whereas salivary analysis may reflect secretory activity from salivary glands, lymphatic system, or brain^22^. Due to their broader dynamic range relative to plasma, urine, and saliva are particularly amenable for deep proteomics profiling utilizing contemporary mass spectrometry (MS)-based proteomic technology and advanced sample preparation methodologies^23^. Simultaneously profiling these body fluids with state-of-the-art proteomics can uncover systemic interaction networks that explain key drivers of proteins mediating exercise-induced adaptations. Furthermore, the increasing availability of data from large-scale consortia and cohorts, including tissue/cell-type specificity^24^, protein quantitative trait loci (pQTL)^25^, and genome/protein-wide association studies (GWAS)^26,27^, presents an unprecedented opportunity to establish the roles of exercise-responsive proteins in health and disease.

This study aimed to elucidate the temporal dynamics of proteins across multiple human body fluids in response to acute exercise and characterize their biological importance. Utilizing high-resolution proteomics, we analyzed saliva, urine, and plasma samples, identifying distinct temporal protein clusters associated with taste perception, iron metabolism, and synaptic function, respectively. Integrative body fluid network analyses uncovered the extracellular matrix and immune activation as central components of the proteomic response to exercise. To validate and extend our findings, we replicated the plasma proteomic changes in an independent cohort of older adults who performed aerobic, high-intensity interval training, and resistance exercise, confirming over 1,000 exercise-responsive proteins and revealing variations influenced by sex and exercise modality. Additionally, by integrating genetic data^26,27^, we linked exercise-regulated proteins to health and disease pathways, identifying exerkines with known approved drugs. Collectively, this comprehensive multi-fluid proteomic resource advances our understanding of the systemic molecular responses to acute exercise and offers insights into potential therapeutic strategies.

## Results

### Study design and clinical characteristics of the Discovery cohort

We recruited 25 healthy young-adult men (BMI = 23 ± 1; *V*O_2max_ = 49 ± 8 mL·kg^-1^·min^-1^; age = 29 ± 3 y) in a Discovery study to explore body fluid dynamics to acute exercise (Tables S1A-C). This study involved three visits: (1) an initial visit for *V*O_2max_ and body composition assessment, (2) a second visit for the acute exercise bout and fasted body fluid sampling, and (3) a final visit for body fluid sampling 24 h post-exercise (Figure 1A). The acute exercise bout consisted of 45 min on a cycle ergometer at ∼75% of their maximum heart rate (HRmax) determined by VO_2max_. Additionally, to avoid post-prandial effects, there was no access to food during the recovery phase (0-3 h post-exercise). Intravenous blood and saliva samples were collected pre- as well as ∼0, 0.5, 1, 3, and 24 h post-exercise, while urine was collected at pre-, ∼0, and 24 h post-exercise (i.e., 15 samples/individual).

**Figure 1.**
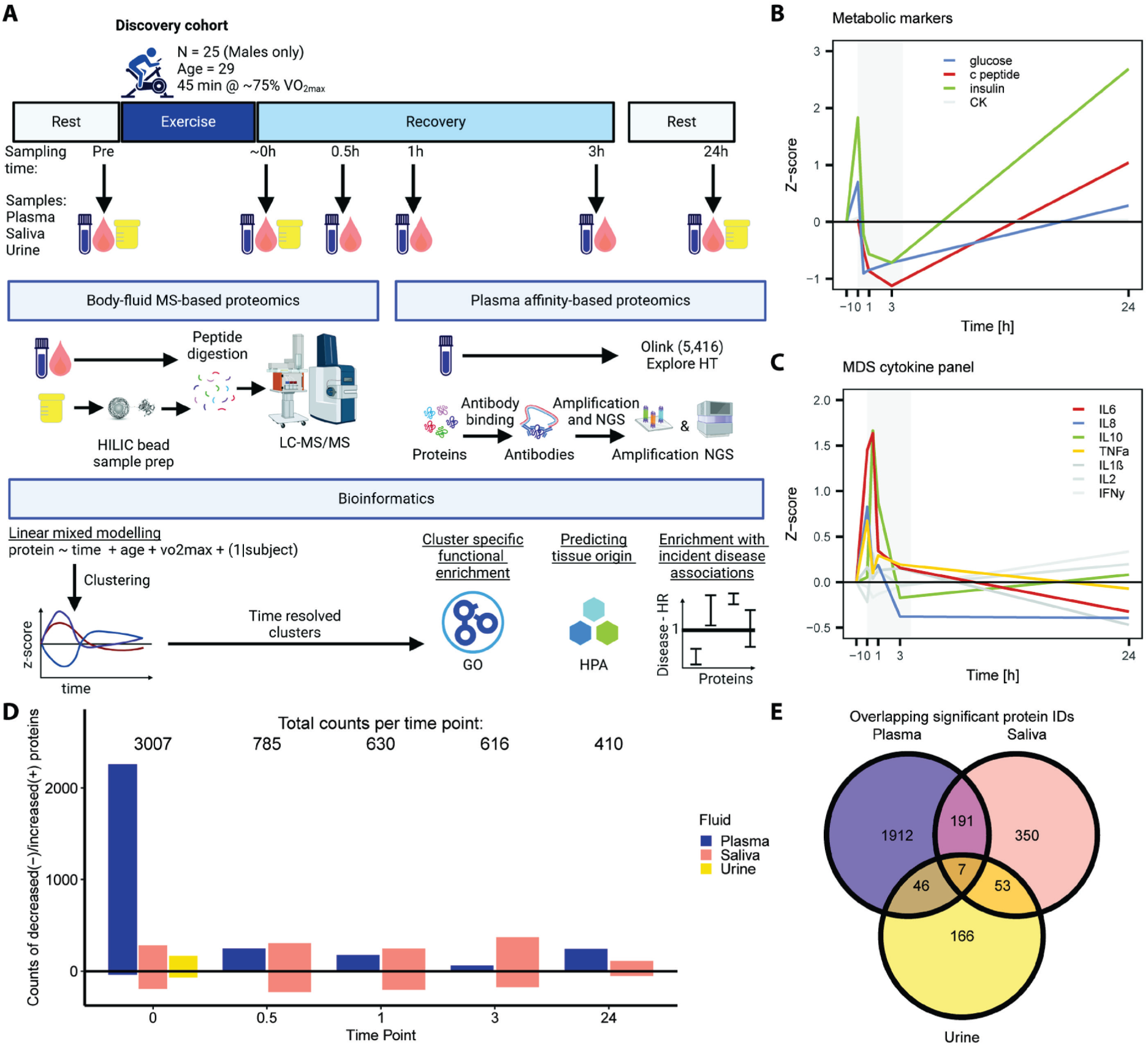
Overview of study design and body fluid proteome responses to acute exercise. **A)** Schematic of the Discovery cohort study design, detailing sampling times, proteomic profiling methods, and bioinformatic approaches created in biorender.com. **B)** Changes in plasma metabolic markers post-exercise relative to pre-exercise levels, including creatine kinase activity (CK U/L), glucose (mmol/L), proinsulin C-peptide (pmol/L), and insulin (pmol/L). **C)** Cytokine response to acute exercise relative to pre-exercise, including IL-6, IL8, IL-10, TNF-α, IL1-β, IL-2, and IFN-γ. Colored lines indicate a significant time effect (p≤0.05); grey lines denote non-significant changes (p>0.05). **D)** Bar plot summarizing the number of significantly regulated proteins at each time point, with total counts displayed above each bar, inclusive of up- and down-regulated proteins. **E)** Venn diagram illustrating overlapping identified and significantly regulated proteins in saliva, urine, and plasma.

First, we measured established metabolic and inflammatory biomarkers in plasma to validate the physiological response to acute aerobic exercise (Figures 1B and 1C; Table S2A). Exercise had a similar effect on glucose (*P* = 7.9e-14), insulin (*P* = 4.2e-03), and c-peptide (*P* = 2.8e-06), showing an initial elevation at 0 h followed by an overcorrection below pre-exercise levels between 0.5-3 h post-exercise (Figure 1B; Table S2A). In the post-exercise period, TNF-α (*P* = 4.2e-06) and IL-8 (*P* = 1.9e-12) increased abruptly, while IL-6 (*P* = 1.7e-18) and IL-10 (*P* = 1.7e-11) exhibited a delayed but more sustained elevation before returning to baseline by 3 h post-exercise (Table S2A). These responses replicated the canonical metabolic and immune responses to acute exercise^28^, affirming the robustness of our acute exercise session.

Using MS and Olink affinity-based proteomics, we measured 10,128 proteins across saliva (19%), urine (27%), and plasma (54%), of which 3,007 (30%) were differentially regulated at 0 h post-exercise compared to pre-exercise across body fluids (Figure 1D; Table S2). Exercise induced an even distribution of up- and down-regulated proteins across the salivary proteome from 0 h post-exercise into recovery (601 proteins/31%). Urinary exhibited a modest change immediately post-exercise (10%/272 proteins), with no proteins being differentially abundant at 24 h post-exercise compared to pre-exercise. In contrast, exercise-induced plasma proteome changes were characterized by a sharp increase in protein abundance immediately post-exercise (37%/2,000 proteins at 0 h), followed by a rapid decline to 75 elevated proteins at 3 h post-exercise. Additionally, 289 proteins were consistently identified and regulated by exercise in plasma, saliva, and urine (Figure 1E; Table S2). This highlights the extensive effect of exercise on proteomic remodeling across multiple body fluids.

### Exercise-induced salivary proteome dynamics have disease-relevant signatures

To explore the dynamic effect of exercise on the salivary proteome, we first examined the degree of molecular individuality across participants. Our analysis revealed a molecular signature for each participant (Figure 2A), underscoring the potential utility of saliva for protein biomarker discovery. Next, we calculated the baseline variance in protein levels across participants and estimated the residual variance not explained by time, without adjusting for additional covariates (Figure 2B). Despite substantial inter-individual variability at baseline, exceeding a 2.7-fold difference for many proteins (Figures 2A and 2B), over 20 displayed consistent temporal responses across participants. Notably, proteins such as PRB2 and SERPINF1 showed robust exercise-induced changes (Figures 2B and 2C). These findings highlight a conserved molecular response to exercise despite baseline heterogeneity.

**Figure 2.**
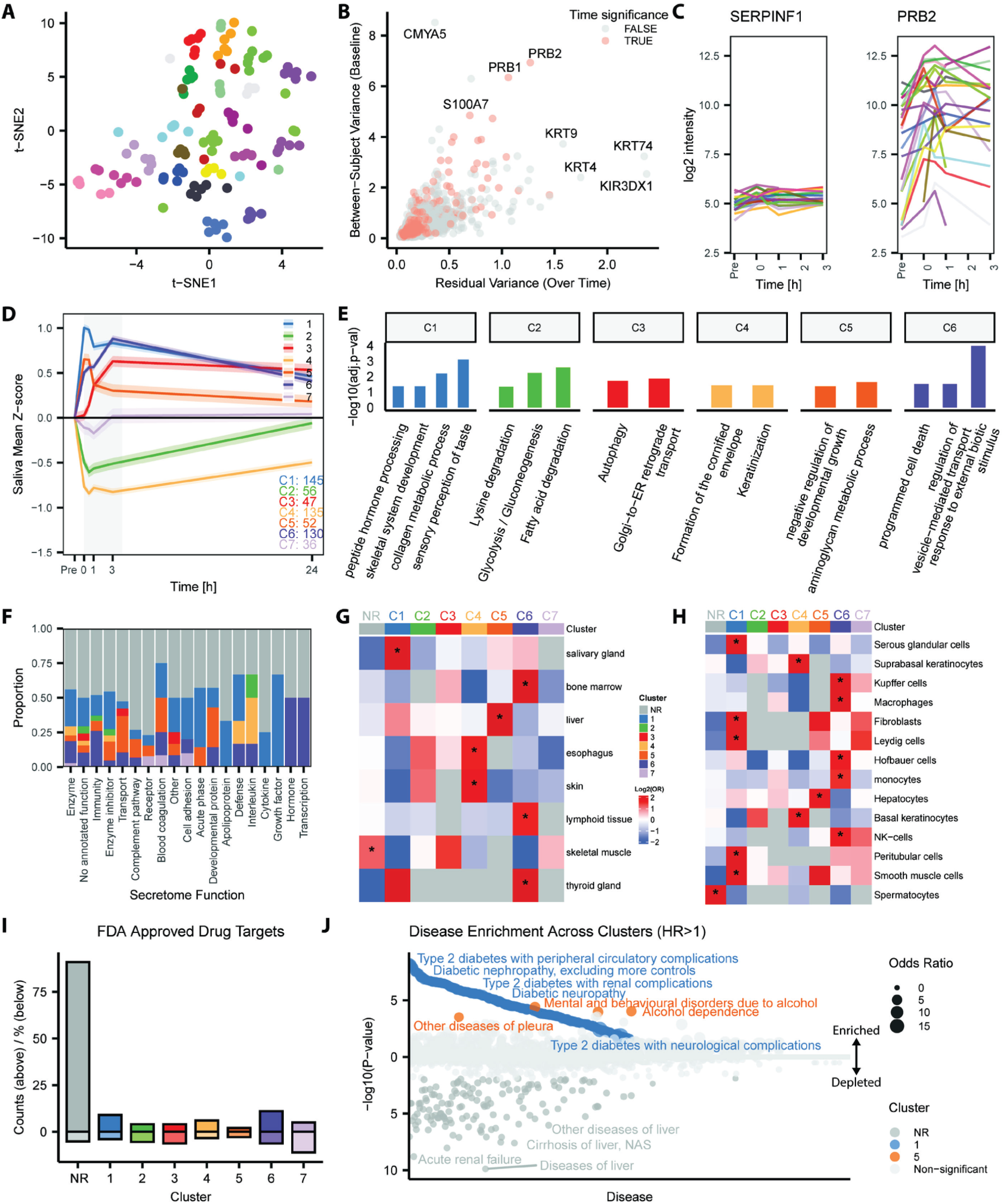
Exercise-induced salivary proteome dynamics have disease-relevant signatures. **A**) t-SNE plot of all samples color-coded by participant ID. **B)** Scatter plot of baseline variance vs. residual variance. **C)** Log2 protein intensities over time, color-coded by participant ID, for proteins with low (SERPINF1) and high (PRB2) baseline variance but low residual variance over time. **D)** Summary of the total number of proteins identified in clusters of protein dynamics following acute exercise (mean ± 95% CI). **E)** Selected overrepresentation analysis results for each cluster identified in panel D. **F)** Proportion of annotated secreted proteins in each cluster. **G)** Tissue and **H)** cell type enrichment of proteins in each cluster. **I)** Counts (top)/percentages (bottom) of identified FDA-approved drug targets. **J)** Incident-disease protein HR>1 enrichment analysis for each cluster. NR = not regulated.

To examine proteomic responses across all body fluids over time, we employed mixed linear models adjusting for age and VO_2max_, while treating participant ID as a fixed variable. In total, 601 proteins, representing 31% (*n* = 1,943) of the salivary proteome, changed in response to acute aerobic exercise (Figure 2D; Table S2B). Next, we performed clustering of temporal patterns of body fluids using a Noise-Augmented von Mises–Fisher Mixture model (NAvMix) and the Bayesian Information Criterion (BIC) for determining the number of clusters^29^. Salivary proteins clustered into six distinct temporal patterns (Figures 2D and S1A-F), while the seventh cluster exhibited stochastic patterns (Figure 2D), highlighted by the protein ECHS1 (Figure S1G). Collectively, these clusters illustrate a robust yet sustained perturbation in the salivary proteome following exercise.

To further interpret the biological relevance of these exercise-induced temporal patterns in the salivary proteome, we performed functional enrichment for each cluster, revealing temporally distinct metabolic shifts in the oral cavity (Figure 2E; Table S3A-B). Specifically, cluster 1, which comprises proteins elevated post-exercise and sustained for 24 h (Figure S1A), was associated with peptide hormone processing and sensory perception of taste (e.g., PIP and AZGP1). In contrast, cluster 2, which contained proteins that initially decreased and then returned to pre-exercise levels by 24 h (Figure S1B), was associated with glucose and lipid metabolism. These findings suggest that acute exercise may modulate eating behavior through dynamic, metabolically linked changes in the salivary proteome.

To uncover the origins of exercise-responsive salivary proteins, we next explored their tissue and cell-type specificity. Using annotation from the Human Protein Atlas (HPA)^24^, we performed cluster-specific enrichment analysis focused on secreted proteins and their putative tissue and cell type specificity^30^. Clusters 1, 5, and 6 were enriched for secreted proteins compared to all other clusters (Figure 2F). These same clusters also showed enrichment for tissue-specific expression in salivary glands (cluster 1), liver (cluster 5), and lymphoid/bone marrow tissues (cluster 6) (Figures 2G and S2). Cell type enrichment analysis indicated that the major contributors to the elevated protein levels were both local (paracrine) and systemic cell-types (endocrine), which included glandular cells, smooth muscle cells, fibroblasts, hepatocytes, natural killer (NK) cells, and macrophage/monocytes (Figure 2H).

Given the link between oral health and the salivary proteome to cardiometabolic disease^31,32^, we investigated whether exercise-regulated salivary proteins were also associated with incident disease. To achieve this, we integrated curated tissue, disease and drug annotations from the DrugBank^33^, HPA^24^, and the newly released protein-incident-disease association atlas from the UK biobank (UKBB)^27^. This approach uncovered 132 FDA-approved drug targets, of which nine were identified within cluster 1 (Figure 2I). Cluster 1 was also highly enriched for metabolic diseases (Figure 2J; Table S4A-B). Specifically, 52/145 unique proteins from cluster 1 have been linked to increased risk of diabetes and its comorbidities (Figure 2J; Table S4). These proteins were functionally related to the extracellular matrix (ECM) (e.g., ELN, PRELP, and SPARCL1), proteolysis/protein processing (e.g., FURIN, CTSB, and AMY2B), lysosomal storage (e.g., TPP1, GLB1, and RNASET2), and general metabolic processes (e.g., NUCB2, IGFBP7, and B4GALT1). These results demonstrate that acute aerobic exercise induces distinct temporal dynamics in the salivary proteome, orchestrated by both local and systemic cellular sources, with many proteins being associated with cardiometabolic diseases.

### The exercise-induced urinary proteome reflects local remodeling and transient shifts in metabolism

Next, we leveraged advances in high-throughput and reproducible urine sample preparation^34^ to investigate the exercise-induced temporal dynamics of the urinary proteome. Urinary proteomic profiles from the same participant showed modest clustering relative to salivary profiles (Figure 3A). Using mixed linear models, we uncovered proteins that had high variability across participants at baseline or across time (Figure 3B). For example, the cytoskeletal protein ACTG1 exhibited higher inter-individual variability at baseline but remained stable over time (Figure 3B). Conversely, signaling proteins like DGKK showed low baseline variance but high variability over time between participants (Figure 3B). Additionally, exercise-regulated proteins, such as PAK4 and MUC5B, displayed variability between participants yet conserved responses over time (Figure 3C). In total, exercise regulated 271 proteins, representing 10% (*n* = 2,767) of the urinary proteome (Figure 3D; Table S2C). These proteins fell into four distinct clusters, largely reflecting a transient shift in urinary proteome composition immediately post-exercise that recovered after 24 h.

**Figure 3.**
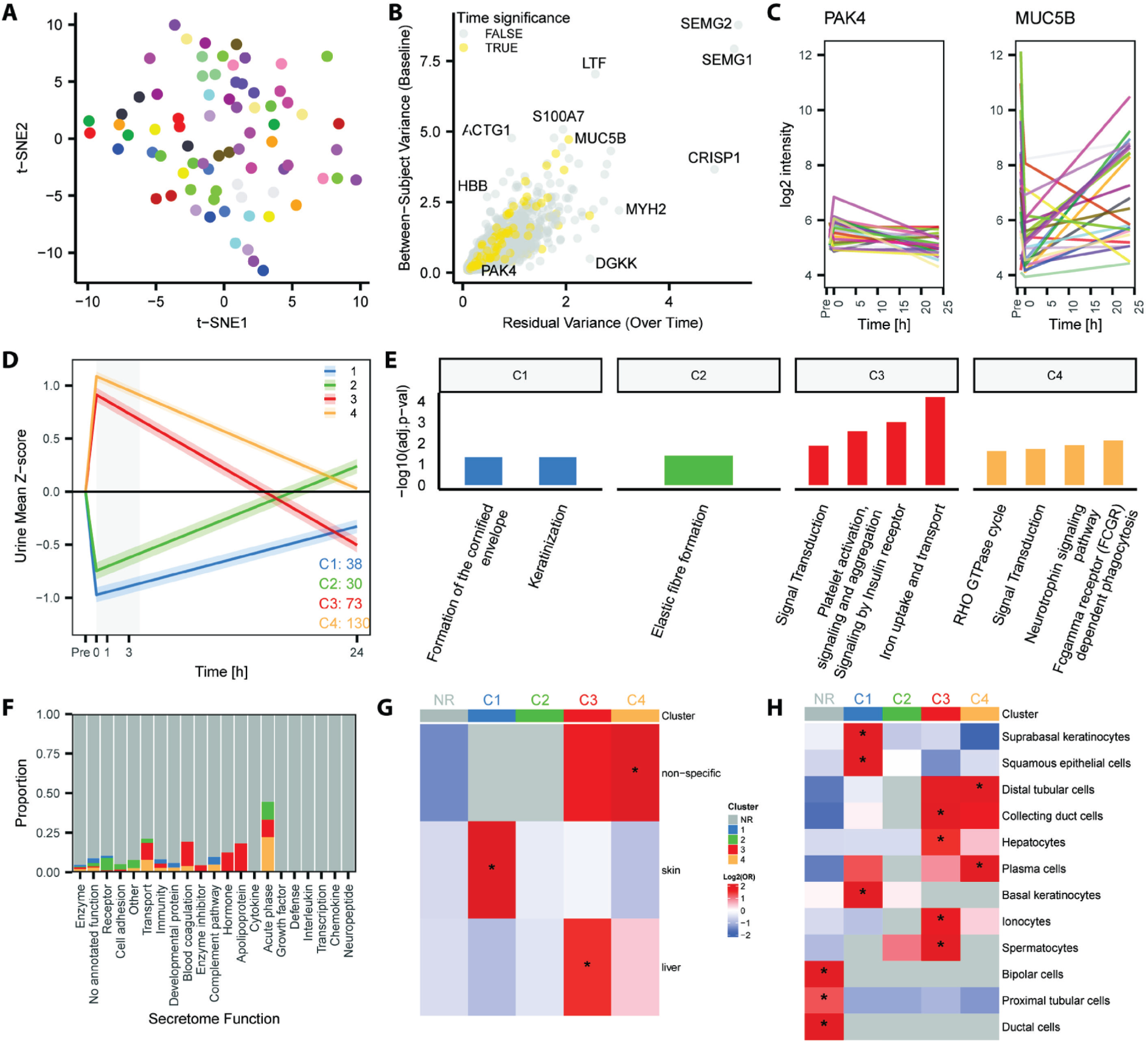
The exercise-induced urinary proteome reflects local remodeling and transient shifts in metabolism. **A**) t-SNE plot of all samples, color-coded by participant ID. **B)** Scatter plot of baseline variance vs. residual variance. **C)** Log2 protein intensities over time, color-coded by participant ID, for proteins with low (PAK4) and high (MUC5B) baseline variance but low residual variance over time. **D)** Summary of the total number of proteins identified in clusters of protein dynamics following acute exercise (mean ± 95% CI). **E)** Selected overrepresentation analysis results for each cluster identified in panel D. **F)** Proportion of annotated secreted proteins in each cluster. **G)** Tissue and **H)** cell type enrichment of proteins in each cluster.

To elucidate biological insights into exercise-induced proteomic changes, we performed cluster-specific functional enrichment analysis (Figure 3E; Tables S3C-D). Clusters 1 and 2, which include proteins that decreased post-exercise and returned to baseline levels by 24 h (Figure S3A-B), were associated with keratinization and elastic fibre formation. Conversely, clusters 3 and 4, which contained proteins that increased post-exercise and returned to baseline by 24 h (Figures S3C-D), were associated with signal transduction, platelet activation, and iron uptake. Additionally, exercise-regulated proteins were depleted for proteins annotated as secreted (Figure 3F). Proteins increasing in abundance were enriched for expression in hepatocytes, collecting duct cells, distal tubular cells, ionocytes, and plasma cells (Figures 3G, 3H, and S4A-C). Conversely, proteins decreasing post-exercise were linked to originating from epithelial cells (Figures 3G, 3H, and S4A-C). In contrast, proximal tubular cells were enriched in the non-regulated proteins, suggesting a segment-specific renal response to acute exercise. These changes may result from shifts in kidney activity/damage, epithelial cell secretion, or preferential uptake of secreted proteins in peripheral tissues. Disease enrichment analysis revealed that while the urine proteome was enriched for kidney disease-protein associations (Figure S3H), these proteins tended not to be regulated by exercise (Figure S3H). Together, these results underscore the kidneys’ role in filtering and excreting byproducts of exercise-induced cellular activity, which may reflect known physiological responses to exercise, including alterations in sympathetic activity, thermodynamics, blood flow, glomerular permeability, or tubular reabsorption.

### The exercise-induced plasma proteome reflects multi-tissue origins and is linked to disease associations

Plasma serves as a dynamic conduit for inter-organ communication, making it a key medium for capturing systemic molecular responses to acute exercise. While MS enabled effective profiling of the salivary and urinary proteomes, its application to plasma is challenged by the inherent complexity and broad dynamic range of plasma. Consequently, MS-based proteome depth by MS was limited and identified a small number of proteins showing regulation in response to exercise (Figure S7; Table S2E). To overcome these limitations and achieve deeper coverage of low-abundant signaling proteins, we employed the Olink Explore HT panel (5,416 proteins) to characterize plasma proteome dynamics.

Among all body fluids analyzed, plasma exhibited the strongest molecular individuality, clustering predominantly by participant identity regardless of post-exercise time point (Figure 4A). Despite high inter-individual variability at baseline, including hundreds exceeding a 2.7-fold difference, the plasma proteome also exhibited the greatest number of proteins with consistent exercise responses across participants (Figure 4B). For example, both NADK and FOLR3 showed robust and reproducible responses to exercise across participants. In contrast to NADK, FOLR3 showed higher baseline variability across individuals, spanning > 128-fold between the lowest to highest observed abundance (Figure 4C). These findings emphasized the robust effect of exercise, even with pronounced baseline heterogeneity in plasma.

**Figure 4.**
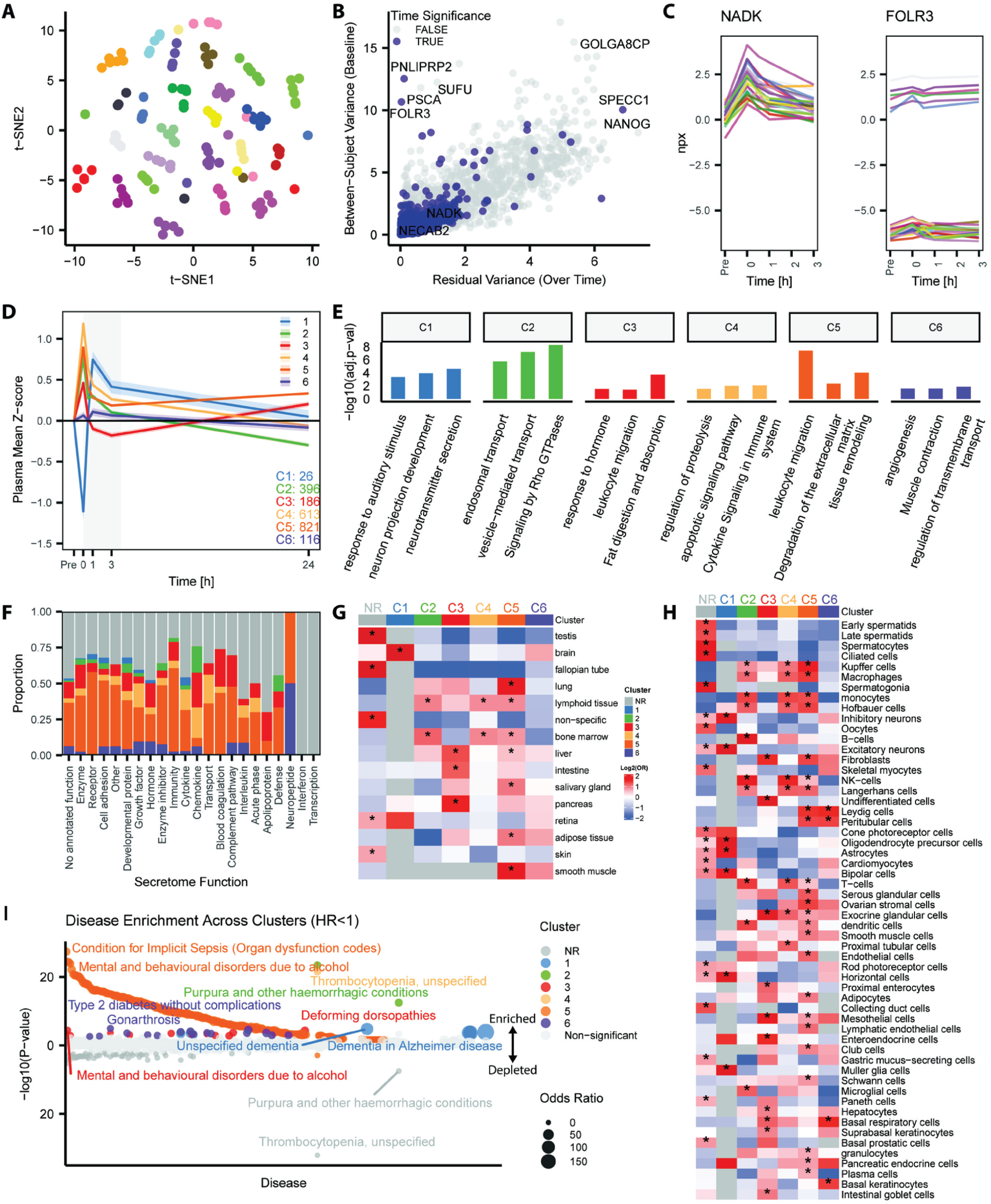
The exercise-induced plasma proteome reflects multi-tissue origins and is linked to disease associations. **A**) t-SNE plot of all samples, color-coded by participant ID. **B)** Scatter plot of baseline variance vs. residual variance. **C)** NPX values over time, color-coded by participant ID, for proteins with low (NADK) and high (FOLR3) baseline variance but low residual variance over time. **D)** Summary of the total number of proteins identified in clusters of protein dynamics following acute exercise (mean ± 95% CI). **E)** Selected overrepresentation analysis results for each cluster identified in panel D. **F)** Proportion of annotated secreted proteins in each cluster. **G)** Tissue and **H)** cell type enrichment of proteins in each cluster. **I)** Protein incident-disease lower risk (HR<1) enrichment analysis for each cluster.

In total, 2,156 proteins, representing 38% of the quantified plasma proteome, were regulated in response to acute aerobic exercise (FDR < 0.05, Figure 2D; Table S2D). These proteins followed five distinct temporal dynamics (Figures 4D and S5A-E), while cluster 6 exhibited stochastic patterns (Figures 4D and S5F). FGF21^35^ and TNFSF11^36^, both in cluster 6 due to their distinct temporal response to acute exercise (Figure S8F), are previously characterized exerkines, thereby supporting the validity of our approach. The high frequency of sampling post-exercise also enabled the identification of small granular shifts in seemingly similar dynamic clusters, like clusters 2, 4, and 5 (Figures S5B, S5D, and S5E). Ultimately, we uncovered a robust and rapid perturbation to the plasma proteome post-exercise.

Functional enrichment of exercise-responsive plasma proteins revealed a large effect on immune and extracellular matrix-related proteins (Figure S5G; Table S3E). Cluster-specific analysis further uncovered shifts in distinct biological processes depending on temporal trajectories (Figure 4E; Table S3F). Specifically, cluster 1, which comprised proteins decreasing post-exercise and exhibited a transient elevation at 1 h post-exercise (Figure S5A), was enriched for synapse functions, including neurotransmitter release (e.g., NPTXR) (Figure S5A). Conversely, cluster 3, which contained proteins like MYOC and POMC (Figure S5C and S8C) that peaked early and exhibited a transient reduction below baseline by 1-3 h, was associated with hormone activity and lipid digestion (Figure S5C). Additionally, cluster 5, the largest cluster, was enriched for secreted proteins (Figure 4F), with many proteins involved in leukocyte function (e.g., SASH3) and tissue remodeling (e.g., CTSB) (Figures S5E and 4E). Tissue and cell-type enrichment analyses revealed broad systemic origins of plasma proteins regulated by exercise (Figure 4G and 4H). Specifically, cluster 1 was linked with brain-specific cell types, cluster 3 with endocrine tissues, like the intestine and pancreas, and clusters 2, 4, and 5 with immune cells. Together, these findings point towards a coordinated, multi-organ proteomic remodeling in response to acute exercise (Figures 4G and 4H).

We next investigated whether exercise-regulated plasma proteins are associated with incident disease^27^. Proteins in cluster 5 formed a signature broadly linked to reduced risk across multiple disease categories, while clusters 1 and 6 were selectively associated with a lower risk for dementia-related diseases and type 2 diabetes, respectively (Figure 4I; Tables S4F-G). For instance, NPTXR, OMG, and VGF, cluster 1 proteins involved in synaptic plasticity, are associated with reduced dementia risk. Twelve proteins in cluster 6 are linked to lower type 2 diabetes risk and annotated for roles in metabolism and neuronal development (e.g., FGF19 and APLP1). Additionally, 16 proteins from cluster 3 are linked to resilience for mental disorders and function in immune and developmental processes (e.g., TNFSF10 and SLITRK6) (Figure 4I). Together, acute exercise elicits plasma proteome changes that are temporally coordinated, biologically diverse, and tissue-specific, which may account for the protective effects of regular exercise training against the manifestation of future diseases.

### Acute exercise coordinates protein networks across body fluids

To capture systemic coordination across the body fluid compartments, we performed weighted protein correlation network analyses^37^ on the proteome data from saliva, urine, and plasma at pre-, 0 h, and 24 h post-exercise, which is described in detail in our Methods section. This framework enabled the identification of co-regulated protein modules and their associations with biological processes, thus revealing dynamic exercise-responsive regulatory networks. Specifically, we observed a doubling in the average number of highly connected proteins (i.e., hub proteins) across body fluids immediately post-exercise compared to pre-exercise and 24 h post-exercise (Figures 5A-B), indicating transient network expansion. This expansion was also accompanied by a change in the top hub proteins (Figure 5C), reflected by the redistribution of proteins from unassigned modules at pre-exercise (grey) into co-expression clusters post-exercise (colors), which largely returned to baseline by 24 h post-exercise (Figure 5D). Overrepresentation analysis of the unique top 500 hub proteins at each time point (Figure 5C; Table S3G) revealed a shift from proteasome complex and protein folding processes at pre-exercise to humoral immune response and the extracellular matrix (ECM) immediately post-exercise (Figure 5E). In contrast, the proteins identified 24 h post-exercise were associated with metabolism and post-transcriptional regulation (Figure 5E), suggesting that immune activation and ECM perturbation initiate the response, while metabolic adaptations predominate the recovery.

**Figure 5.**
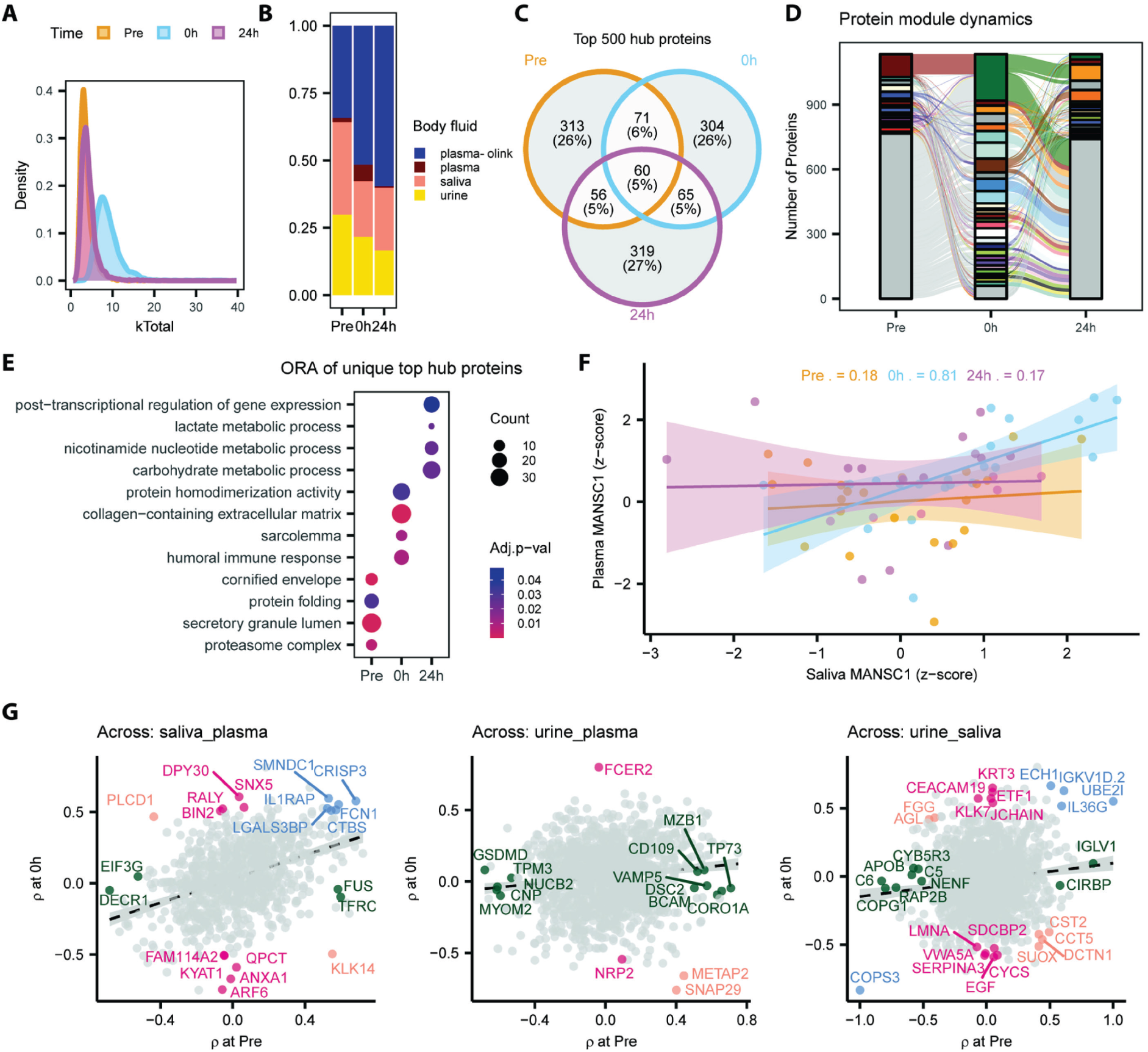
Acute exercise coordinates protein networks across body fluids. **A**) Number of edges per protein, indicating protein-protein correlations from WGCNA. **B)** The proportion of each body fluid to the WGCNA network. **C)** Overlap in the top 500 hub (most edges) proteins from WGCNA. **D)** Protein module dynamics across time points: each color indicates a unique network module at each time point. **E)** Overrepresentation analysis from unique hub proteins at each time point. **F)** Example of time point-specific correlation of a protein across body fluids (MANSC1). **G)** Scatter plots of correlation of proteins identified across body fluids at pre- (x-axis) and 0 h post-exercise (y-axis) (blue = correlated at both pre-exercise and 0 h post-exercise, pink = correlated at 0 h but not pre-exercise, green = correlated at pre-exercise but not 0 h post-exercise, and orange = shifting the direction of correlation). WGCNA = Weighted Correlation Network Analysis; ORA = Overrepresentation Analysis.

To further investigate inter-fluid coordination, we focused on proteins detected across the three body fluids and generated a time-resolved correlation matrix. Several proteins exhibited exercise-dependent associations across compartments. For example, MANSC1 levels in saliva and plasma showed no correlation at pre-exercise or 24 h post-exercise but became strongly correlated immediately post-exercise (Figure 5F). This points towards a system-wide perturbation in MANSC1 following exercise that is due to either an increased mobilization from plasma or a conserved body fluid-independent response. We further expanded on these examples and uncovered that saliva and plasma have more correlating proteins independent of time compared to comparisons with urine (Figure 5G). We identified proteins with distinct association patterns: those that remained consistently associated regardless of time (blue), those whose associations were lost following exercise (green), those that became associated only after exercise (pink), and those that exhibited a complete reversal in the direction of association (orange). One example was SNAP29, a SNARE protein involved in vesicle trafficking^38^. Specifically, its correlation between plasma and urine shifted from positive at pre-exercise to negative immediately post-exercise (Figure 5G, 2^nd^ panel), potentially suggesting reduced excretion or systemic sequestration. Additional proteins that reduce their correlation across plasma-urine post-exercise were associated with cytoskeletal organization (e.g., TPM3 and CORO1A), cell adhesion (e.g., DSC2 and BCAM), and immune system functions (e.g., GSDMD, MZB1, and CORO1A). Conversely, several proteins in urine and saliva remained negatively associated across time, like COPS3, which regulates protein degradation, whereas other proteins started to become positively or negatively correlated, like the membrane protein CEACAM19 and growth factor EGF. In summary, these analyses reveal that acute exercise induces transient reorganization of protein co-expression networks across body fluids, with inflammation and ECM remodeling as an early response, followed by metabolic realignment during recovery. Additionally, a coordinated shift in inter-fluid protein relationships suggests selective routing, retention or excretion of specific proteins in response to physiological demand.

### Replication of the exercise-induced plasma proteome across age, sex and exercise modalities

Recognizing that exercise encompasses diverse modalities and that protein dynamics following acute exercise are influenced by age and sex, we aimed to replicate the identification of 2,156 exercise-regulated proteins in plasma in a separate cohort. The Replication cohort comprised 24 middle-aged adults (55±8 years), including both men and women (15/24 women), and a randomized crossover design involving three time-matched exercise modalities: aerobic, high-intensity interval training (HIIT), and resistance exercise (Figure 6A). Of the 2,156 exercise-responsive proteins identified in young males, 1,161 were replicated, 358 were newly identified, and while only 2 proteins had an exercise mode by time interaction, 590 had a main effect for exercise modality indicating differences in effect sizes post-exercise across modalities (Figure 6B; Table S2F-H). Comparing the β coefficients and corresponding ratios for overlapping significant proteins (1,161 proteins) between cohorts revealed a predominantly concordant direction of responses, as indicated by a shift in the density distributions toward positive ratios. At 0 h post-exercise, 87% of proteins exhibited a positive beta ratio, indicating concordance between studies (Figure 6C). This concordance decreased over time, with 54% of proteins exhibiting a positive ratio at 0.5 h and 34% at 24 h post-exercise. Notably, the proportion of proteins with extreme beta ratios (absolute value > 2) increased from 2% at 0 h to 14% at 0.5 h, reaching 45% and a bimodal distribution at 24 h, reflecting variability in the protein response to acute exercise during the recovery phase across cohorts (Figure 6C). These findings suggest that while there are conserved exercise-induced proteins, a complex interplay between age, sex, and exercise modality collectively shapes the acute exercise-induced plasma proteome.

**Figure 6.**
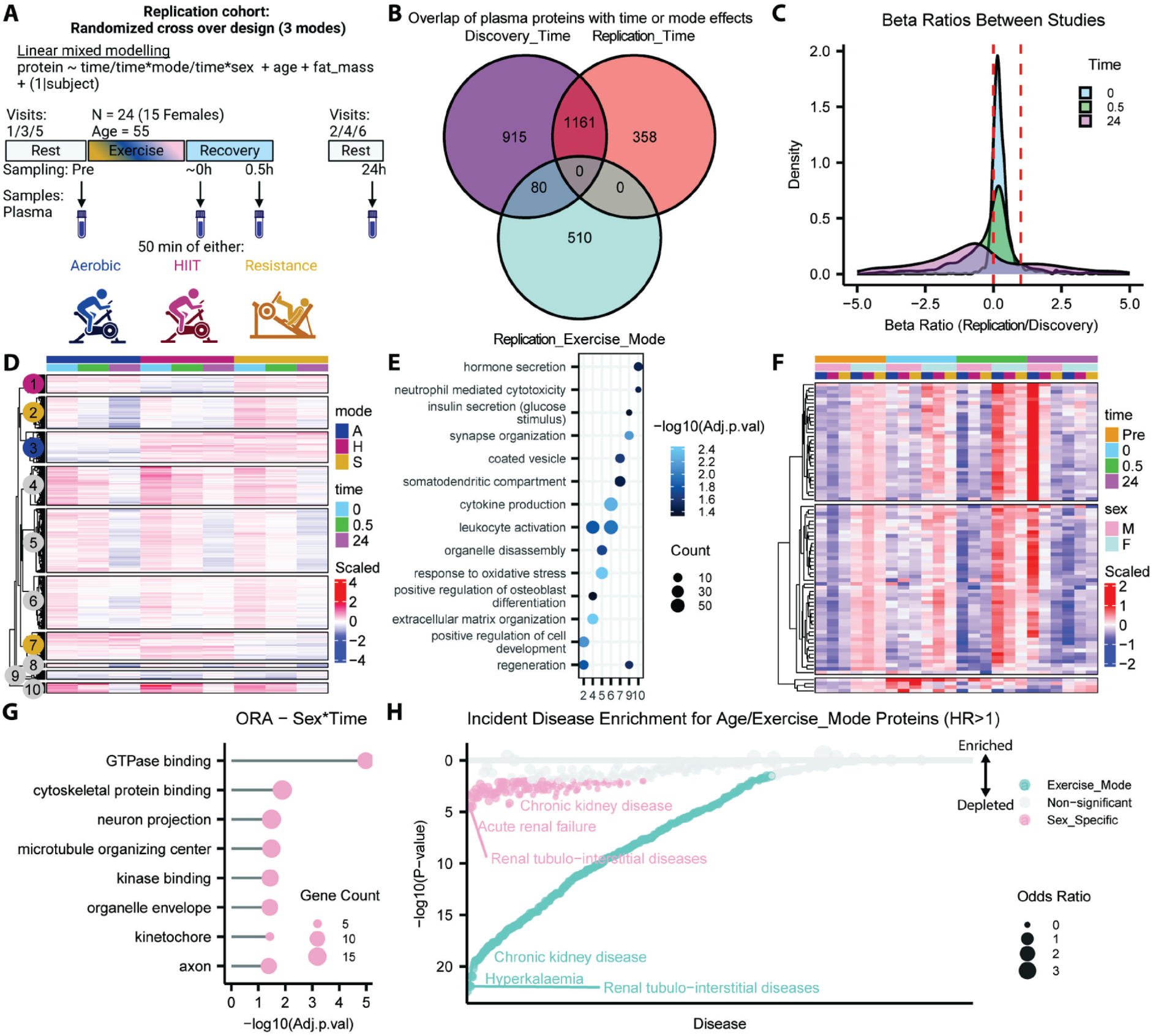
Replication of the exercise-induced plasma proteome across age, sex and exercise modalities. **A**) Schematic representation of the Replication cohort study design created in biorender.com. **B)** Venn diagram illustrating the overlap between plasma proteins with significant time effects in the Discovery cohort and those with significant time and exercise mode effects in the Replication cohort. **C)** Density plot of beta ratios for the Discovery and Replication cohorts for each time point post-exercise. **D)** Heatmap displaying Beta coefficients relative to pre-exercise across different exercise modalities. **E)** Overrepresentation analysis of exercise mode-specific protein clusters identified in panel D. **F)** Heatmap of scaled NPX values across time points for proteins exhibiting significant sex-by-time interactions. **G)** Overrepresentation analysis for proteins with significant sex-by-time interactions from panel F. **H)** Protein incident-disease risk (HR>1) enrichment analysis for proteins identified as exercise mode- or sex-specific. HR = Hazard Ratio; ORA = Overrepresentation Analysis.

To disentangle modality-specific effects, we plotted the scaled effect sizes across all proteins, emphasizing that most exercise-responsive proteins were regulated independently of modality and the main differences were in the magnitude of effect (Figure 6D; Table S2G). However, specific clusters, namely cluster 1, 2, 3, and 7, exhibited divergent responses depending on the exercise mode (Figure 6D). An overrepresentation analysis highlighted that clusters 2 and 7, which showed opposing effects in resistance exercise, were enriched for cell development (higher effect) and coated vesicle cellular component (lower effect), respectively (Figure 6E; Table S3H). Independent of exercise mode, 83 proteins exhibited sex-specific responses to acute exercise (Figure 6F; Table S2H). These proteins were enriched for cellular components, such as axon, organelle development, and cytoskeletal terms (Figure 6G; Table S3I). Disease enrichment analysis also revealed that relative to the background proteome, proteins influenced by exercise mode or sex were depleted for associations with incident disease (Figure 6H; Table S4G-H). Collectively, these findings suggest that while differences exist between the plasma proteome response across age, sex, and exercise mode, the proteins exhibiting consistent regulation are more strongly linked to disease associations and therefore likely to contribute to the long-term health benefits of exercise.

### Genetics and pharmacological integration link exercise-regulated proteins to disease traits

To investigate the potential relevance of exercise-regulated proteins to their long-term health benefits, we examined their genetic associations with health and disease-related traits. We performed colocalization analyses between exercise-regulated proteins and traits modifiable by physical activity, focusing on cis-acting protein, expression, and splicing quantitative trait loci (pQTLs, eQTLs, sQTLs) located within 1 Mb of the corresponding protein-coding genes (Figure 7A; Table S5). This framework leverages genetic evidence for linking changes in protein expression and health and disease traits, enabling the elucidation of potentially causal pathways. In general, most exercise-regulated proteins have evidence for changes in their expression being linked to health and disease traits, as indicated by the number of edges in the full network (Figures 7B and 7C). However, proteins showing the strongest fold changes, increasing and decreasing, tended to have fewer trait colocalizations (Figure 7B), suggesting a potential trade-off between the amplitude of response and genetic evidence for pleiotropy. To further explore their biological relevance, we mapped these proteins to their known drug targets using DrugBank (v5.1.13)^33^ (Figures 7A and 7C). The resulting integrative network, comprising over 750 proteins, 250 traits, and 250 drugs, revealed four distinct sub-communities each organized around related trait categories: (1) physical activity and function, (2) endocrine and digestive health, (3) cardiovascular health, and (4) neurological and body composition traits (Figure 7C). Notably, metabolic traits span all communities and hold a central position in the network, underscoring metabolism as a central axis of exercise adaptation and disease prevention (Figure 7C).

**Figure 7.**
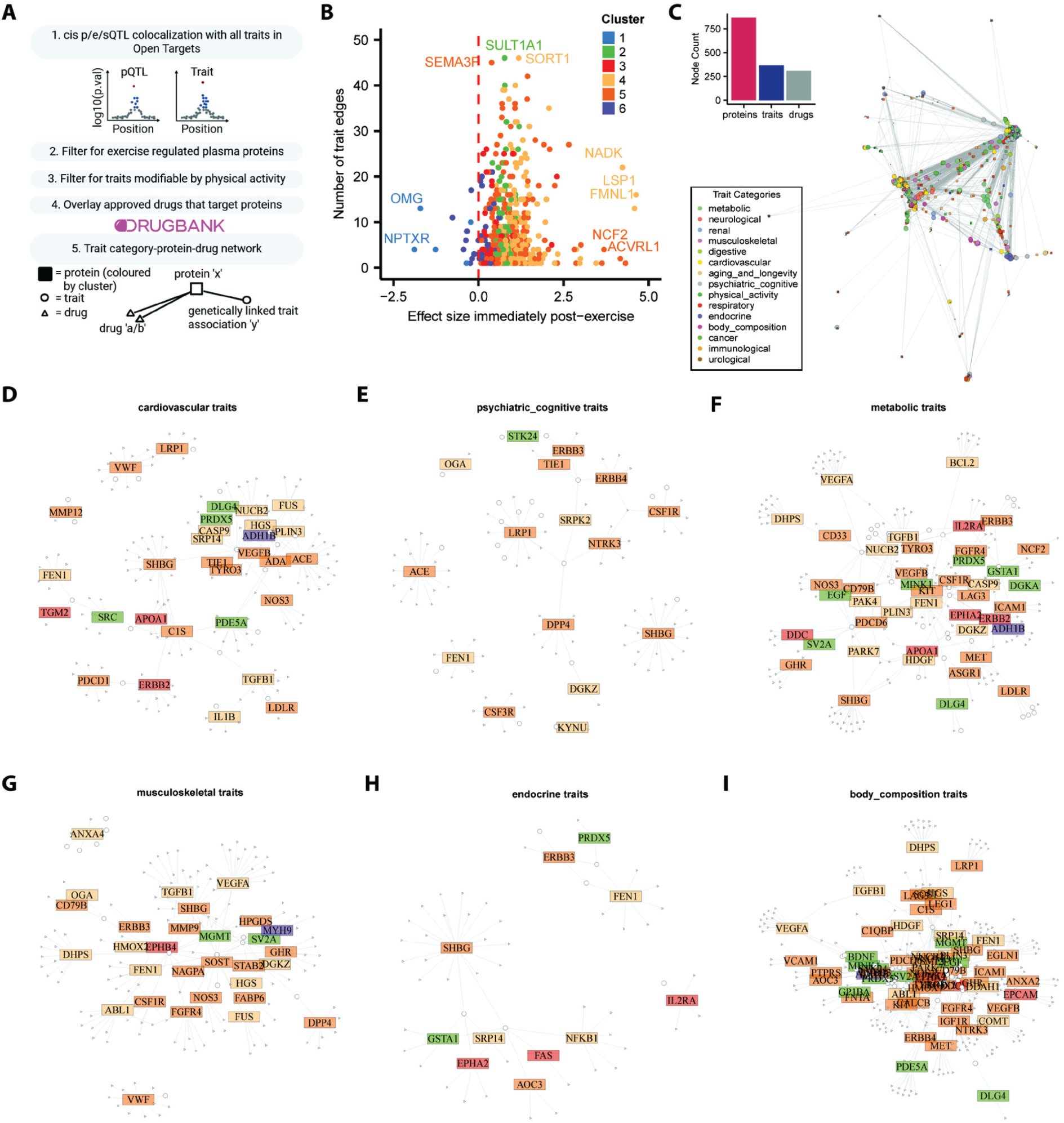
Genetics and pharmacological integration link exercise-regulated proteins to disease traits. **A**) Schematic workflow illustrating the colocalization analysis process used to construct the exercise-inducible protein-trait-drug network created in biorender.com. **B)** Scatter plot of effect sizes for pre- to 0 h post-exercise (x-axis) and the number of significant cis p/e/sQTL-physical activity modifiable trait edges for exercise-regulated proteins. **C)** Comprehensive network diagram showing proteins (squares) color-coded by cluster, with hubs representing specific trait categories labelled accordingly. **D-I)** Subnetworks of specific trait categories (defined in panel titles and white circles in sub-networks) that have protein-drug edges (triangles). p/e/sQTL = protein/expression/splice quantitative trait loci.

We next explored these sub-communities that had known approved drug targets. These networks highlight that exercise might regulate several drug targets, indicating it influences diverse pathways important for disease management (Figures 7D-I). For example, DPP4 inhibitors, widely used and now generic for the treatment of diabetes^39^, also show associations with psychiatric and cognitive traits (Figure 7E). Similarly, levetiracetam, an anticonvulsant for epilepsy targeting SV2A, is linked to metabolic and body composition traits (Figures 7F and 7I). These observations suggest that exercise exerts its effects on pathways important for disease management and that several approved drugs may have unknown pharmacodynamics and pleiotropic effects that should be explored for new (contra)indications (Figure 7A). It also reinforces that the molecular complexity of acute exercise is unlikely to be fully recapitulated by any single compound or a drug combination.

## Discussion

This study presents a comprehensive, time-resolved characterization of the acute exercise-induced multi-compartmental human proteome, capturing granular temporal dynamics across plasma, saliva and urine. By integrating deep-proteomic profiling with network analysis, tissue-specific protein annotations (HPA^24^), disease associations^27^, genetics data (Open Targets^26^), pharmacological mapping^33^, and cross-cohort replication, we identified coordinated and tissue-linked proteomic responses to exercise. These molecular signatures are broadly associated with incident disease and traits that are modifiable through physical activity. Our findings advance our molecular understanding of how exercise remodels the human body and provide a foundational resource to explore candidate exerkines as potential biomarkers or therapeutic targets to improve human health.

Our analysis extends the findings from previous seminal papers^14,15,18,40^ by elucidating and characterizing dynamic proteome dynamics across multiple body fluids. Plasma proteins exhibited pronounced elevations, predominantly returning to baseline within 1 h, consistent with prior investigations^14,15,40^. We identified distinct temporal profiles of proteins, including hormone-like responses in plasma proteins originating from the pancreas and intestine, and sigmoidal patterns associated with synapse function and future dementia risk. In contrast to plasma, salivary proteins demonstrated a balanced distribution of up- and down-regulation, maintaining consistent alterations from immediately post-exercise throughout the recovery period, suggesting distinct regulatory mechanisms. Notably, salivary proteins exhibiting prolonged elevation post-exercise were implicated in taste perception and broadly associated with cardiometabolic diseases, including type 2 diabetes. These findings align with our recent findings identifying a salivary proteomic signature that distinguishes individuals with type 2 diabetes from healthy controls^31^, indicating that the salivary proteome responds to both metabolic disease and acute exercise. The urinary proteome exhibited alterations that resolved within 24 h post-exercise, consistent with previous evidence^21,41^ linking acute exercise to metabolic perturbations in urine. These compartment-specific differences likely reflect underlying physiological and anatomical factors, including autonomic nervous system activity, fluid flow dynamics, and tissue-specific protein leakage. Tissue enrichment analysis substantiated these interpretations, which included evidence for exercise-regulated plasma proteins being derived from diverse organs, urinary proteins predominantly originating from the kidney and liver, and salivary proteins originating from glandular and immune tissues. Collectively, these results emphasize the heterogeneous biological inputs driving exercise-induced responses across body fluid compartments and demonstrate the utility of multi-fluid profiling for identifying fluid-specific exerkines. Along with molecular insight, our temporally resolved tissue-specific protein clusters have the potential to serve as targeted panels that warrant further investigation as potential biomarkers for adaptation in both health and disease states.

Despite distinct proteomic composition and dynamic profiles across body fluids, our network analysis revealed coordinated proteome remodeling following exercise. This integrated response was characterized by enhanced interconnectivity across fluid compartments and encompassed proteins associated with ECM remodeling, immune activation, and protein homodimerization. While previous investigations have demonstrated robust ECM^40^ and immune system^14^ responses to acute exercise, our findings extend this knowledge by identifying ECM-related proteins as key network hubs with high connectivity and functional diversity. These included structural proteins (e.g., COL6A1/2, SPARC, and DAG1), components of vesicle transport (e.g., APOA1, CLU, and AZGP1), and modulators of cell signaling (e.g., ANGPTL2, AEBP1, and PRG2). Concurrently, proteins enriched for homodimerization activity comprised highly connected nodes involved in EV biogenesis/cargo (e.g., TSG101, TPD52, and APOA1), cytoskeletal organization (SLK, CACYBP, and PTPA), and signal transduction (STAT5B, IDE, BCL2L1, and ERN1). Consequently, we postulate that alterations in ECM composition and stoichiometry in protein homodimerization may facilitate the mobilization, functional processing, and targeted delivery of exerkines following exercise. This hypothesis is corroborated by previous studies demonstrating integrin-mediated tissue-specific uptake of extracellular vesicles in both cancer^42^ and exercise^18^. Further mechanistic investigations into these pathways may elucidate how exercise orchestrates tissue crosstalk and systemic adaptation at the molecular level.

Our replication analysis in middle-aged adults confirmed numerous exercise-induced plasma proteomic alterations observed in the initial Discovery cohort and identified new proteins influenced by exercise mode and sex. Notably, while subtle differences were evident between the plasma proteome responses across age, sex, and exercise mode, the proteins exhibiting consistent responses demonstrated enrichment for associations with incident disease, particularly cardiometabolic and neurological disorders^27^. This observation suggests that the most consistently exercise-responsive proteins across cohorts may serve as the core mediators of the long-term health benefits of exercise. These findings corroborate previous investigations linking plasma proteomic signatures to cardiorespiratory fitness improvements following exercise training^43^ and support the hypothesis that the protective effects of exercise are predominantly conserved across diverse modalities and sex differences. Indeed, even brief and intermittent bouts of exercise enhance cardiorespiratory fitness and attenuate disease risk^44^, reinforcing the concept that the molecular pathway engagement, rather than the specific exercise modality, primarily drives systemic adaptive responses.

Integration of genetic evidence with exercise-responsive plasma proteins revealed potential pathways that mediate long-term health benefits. Several exercise-regulated plasma proteins demonstrated genetic associations with traits that are modifiable by physical activity, including type 2 diabetes (e.g., NUCB2), blood lipids (e.g., NPTXR), anthropometric parameters (e.g., MANSC1), tissue monocyte counts (e.g., SNAP29), and cognition (e.g., MYOC and IST1). Some of these associations are ubiquitously identified across tissues, like SNAP29, which we found to be sequestered in plasma post-exercise, while others exhibit tissue-specific relationships (e.g., NPTXR and brain tissue). These findings suggest that transient alterations in protein exposure following acute exercise originate from diverse tissues and may contribute to distinct long-term physiological adaptations. This interpretation aligns with previous research demonstrating that exercise-induced tissue-specific gene expression modifies disease risk profiles^45^. Despite the considerable molecular complexity of the exercise response, our analysis identified >200 exercise-regulated proteins that are known targets of FDA-approved pharmacological agents, highlighting the robust effect of disease-modifiable pathways, opportunities for therapeutic repurposing, and the development of new therapeutics that mimic discrete aspects of exercise biology. Nevertheless, the extensive and coordinated exercise-induced proteomic remodeling reinforces that no pharmacological intervention can fully recapitulate the systemic benefits of exercise.

In conclusion, our investigation provides compelling evidence that acute exercise induces distinct, temporally dependent alterations in the body fluid proteomic landscape, reflecting a coordinated, multi-system response. These findings advance the understanding of the molecular drivers of exercise-induced adaptations and establish a foundation for the informed development of diagnostic biomarkers and therapeutic intervention strategies targeting exercise-responsive molecular pathways.

### Limitations of the study

Several methodological constraints warrant consideration when interpreting our findings. Firstly, our experimental design presupposes that post-exercise proteomic alterations are attributable to the exercise intervention. This potentially overlooks proteins subject to circadian oscillations. The limited number of protein-disease associations that were observed with exercise-regulated urinary proteins (Figure S3H) likely reflects restricted coverage of urinary proteomics using LC-MS based proteomics in comparison to the 3K Olink panel employed for protein-disease mapping in the UKBB^27^. Furthermore, urine sampling was restricted to only 3 discrete time points, constraining our network analysis. In our Replication cohort, we exclusively monitored plasma proteomic changes, which prevented our replication of exercise-induced salivary and urinary proteomic alterations. Additionally, our plasma MS-based proteomics approach demonstrated limited coverage, preventing internal validation of our affinity-based proteomic findings. Consequently, these findings should be validated across multiple platforms to account for potential discrepancies arising from technological differences^46–48^. Our colocalization analysis should be considered exploratory and serves primarily as a foundation for subsequent investigations. Specifically, rigorous validation of the temporal dynamics of exercise-regulated proteins and how they mechanistically influence various health and disease phenotypes remains imperative. It is also critical to emphasize that these proteins have pleiotropic roles in health and disease, and despite many exerkines likely contributing to the beneficial effects of exercise, there are also counterregulatory factors identified that, if targeted, could be harmful. Thus, there is a need to complement this data with future resources to disentangle which proteins are appropriate targets for therapeutic lead development or drug repurposing. Finally, given our small and homogenous sample, future large-scale studies are needed, like the Molecular Transducers of Physical Activity Consortium (MoTrPAC)^49^, which will be sufficiently powered to explore some host factors, including how the genetic landscape influences proteome dynamics in response to acute exercise.

## Methods

### Experimental model and study participant details Discovery cohort

#### Participants

25 healthy, normal weight, male participants were included through advertisement on research recruitment webpages and at educational institutions within the Capital Region of Denmark. Inclusion and exclusion criteria can be found in Table S1A. All participants provided written informed consent prior to participation. The study protocol was approved by The Scientific Ethics Committee of the Capital Region of Denmark (Journal number H-16025903) and The Danish Data Protection Agency (Journal number RH-2016-239). The study was performed in accordance with The Helsinki Declaration.

#### Study design

The study consisted of three days: a screening day, a test day, and a post-test day (Figure 1A). During the screening day participants underwent a detailed medical interview and health examination. The latter included blood sampling to screen for chronic conditions such as diabetes, kidney, and liver disease. Anthropometric measurements (height, weight, waist- and hip circumference), a DXA-scan for body composition, and a VO_2max_ test to assess fitness levels were also conducted (Table S1B).

On the test day participants arrived in the morning following an overnight fast (12h), with the allowance to drink water up to two hours before testing. Participants were instructed to refrain from vigorous physical activity and alcohol consumption 24 h prior to testing and were encouraged to use passive means of transportation. The test day began with a 45 min resting period during which participants were placed in a semi-supine position. This was followed by an acute exercise session during which participants biked on an ergometer bike (Ergorace 2 LTD, Kettler) for 45 min at an intensity corresponding to 75% of their heartrate reserve capacity (maximal HR (from the VO_2max_) - resting HR (from the resting period) * 0.75 + resting HR). During the exercise, each participant consumed a fixed volume of 300 ml of water.

The exercise bout was followed by a post-exercise resting period where they refrained from eating, during which blood, urine and saliva samples were collected at the time points illustrated in Figure 1A. The post-test day was dedicated purely to the collection of blood, urine, and saliva samples 24 h after the exercise session. Instructions regarding fasting, alcohol consumption and transportation were consistent with those prior to the test day.

### Replication cohort

#### Participants

24 healthy inactive adult participants (55±8 years and 62.5% female; Table S1B) were included through recruitment within the Capital Region of Denmark (Inclusion and exclusion criteria can be found in Table S1A). Recruitment was conducted through a recruitment agency (e.g. www.ForskningNU.dk). All participants provided written informed consent before participation. The study protocol was approved by the Scientific Ethics Committee of the Capital Region of Denmark (Journal number H-22040452) and the Danish Data Protection Agency (Journal number RH-2017-60). The study was performed in accordance with the Helsinki Declaration

#### Study design

This trial was a single randomized crossover design where participants underwent three sets of exercise sessions. A set consisted of one exercise modality performed twice across 7 weeks, including the baseline measurements. The exercise modalities include 1) continuous aerobic exercise, 2) high-intensity interval exercise and 3) resistance exercise training. A set consists of test/familiarization and retest of the same exercise mode. For this study, we only examined plasma samples from the initial test day for each of the three exercise modes.

#### Randomization, sequence generation, and allocation concealment

Participants were randomly allocated by an independent statistician who used a computer-generated randomization schedule including six possible sequence orders using balanced blocks, stratified by sex. The schedule was forwarded to a secretary not involved in any study procedures and stored on a password-protected computer. Sequentially numbered (according to the sequence) opaque, sealed envelopes were prepared and stored in a locked cabinet in an access-restricted room. The envelopes were lined with aluminum foil to render them impermeable to light. Following the baseline measurements, a study nurse, not involved with any study procedures, opened envelopes, and informed the researcher about the order. The participants were informed about the trial conditions on arrival to the lab in the morning for each exercise session.

#### Exercise modes

The exercise modes were matched based on time, equating to 50 min of exercise per session. They were repeated in sets on consecutive weeks (e.g. three Tuesdays in a row) with a 1-week wash-out period. The trial conditions included three exercise modes described below:

#### Continuous aerobic exercise session

The session began with a 10 min warm-up, followed by 40 min of continuous aerobic exercise at an individualized intensity of 64-76% of their maximal heart rate (HRmax) on a cycle ergometer (Kettler).

#### High-intensity interval training (HIIT) session

The session began with a 10 min warm-up followed by 25 min of HIIT (5 bouts of 4 min at >85% HRmax interspaced by 4 min of low intensity training) and a 4 min cool-down on a cycle ergometer (Kettler).

#### Resistance exercise session

The session began with a 10 min body weight-based dynamic warm-up, including five exercises with three sets each with a 15 s/15 s work/rest ratio. Subsequently, participants completed five sets of four machine-based exercises: leg press, chest press, leg extension, and seated row. Each exercise began with two warm-up sets at 30% and 50% of the individual’s one-repetition maximum (1RM), followed by three working sets at 70% of 1RM. Each set consisted of 10 repetitions, with a 2 min rest interval between sets, totaling approximately 2.5 min per set.

## Method details

### Sample collection

Blood samples were collected from an antecubital vein in EDTA tubes, which were immediately placed on ice or in a cooled centrifuge (4°C). Blood was spun at 2000 x *g* for 10 min followed by pipetting of the plasma into 1.5 mL microcentrifuge tubes. Prior to saliva sampling, the participants rinsed their mouths with sterilized sodium chloride (3 x 30 seconds) followed by 5 min of rest. Saliva production was stimulated by chewing a sterilized cotton swab (Salivette) for one minute at a frequency of 1 chew/s. The cotton swab was collected into a tube and centrifuged at 4°C and 1000 x *g* for 2 min. Urine samples were collected by the subject in a sterile cup, followed by centrifugation for 10 min at 3500 x *g* and 4°C. The supernatant was collected and stored at -80° C until further analyses were performed.

Screening day blood samples were analyzed in accordance with standard clinical practice at the Department of Clinical Biochemistry, Rigshospitalet. Glucose, C-peptide, insulin, and creatine kinase analyses were performed as batch analyses at the Department of Clinical Biochemistry, Rigshospitalet. Glucose and creatine kinase levels were determined by enzymatic absorption photometry (Cobas 8000, module c702 and Cobas Pro, c503 module respectively), whereas C-peptide and insulin levels were determined by Sandwich Electrochemiluminescence-immunoassay (Cobas 8000, module e801). For insulin analyses, a hemolysis index < 50 was accepted. Cytokines were analyzed using the Meso scale Discovery (MSD) V-PLEX Custom Human Biomarkers kit in accordance with the manufacturers protocol.

### LC-MS proteomic saliva sample preparation

Saliva proteomic sample preparation was performed using an in-solution protocol optimized for saliva. Saliva samples were thawed at 4°C and then vortexed. A volume of 20 μL was transferred to a 96-well MS plate containing 20 μL of lysis buffer (1% SDC, 50 mM Tris pH = 8.5). Samples were heated for 10 min at 99°C, cooled to room temperature, then sonicated for 15 min in a Bioruptor water bath sonicator. Approximately 30 μg proteins were digested overnight at 37°C (750 rpm) with Trypsin (1:100 = 0.3 μg) and Lys-C (1:500 = 0.06 μg). Digestion was stopped by acidification to 1% TFA, followed by centrifugation at 4000 x *g* for 10 min to pellet SDC precipitate. Supernatant-containing peptides were desalted using 3xlayer C18 discs plugged into a p200 tip. Samples were eluted with 50 µL of 40% acetonitrile (ACN). Peptides were dried completely in a vacuum concentrator (Eppendorf) and stored at -20°C until further analysis.

### LC-MS proteomic urine sample preparation

Urine proteomic sample preparation was performed using a HILIC solid-phase extraction protocol^34^. Samples were thawed at 4°C and mixed with 300 μL of a modified solubilization buffer (8M Urea, 2% SDS, 50 mM Tris pH=8.5, 10 mM TCEP, 40 mM CAA) in deep-well KingFisher plates. The plate was sealed and placed in a thermomixer at 30°C for 30 min at 800 rpm. The Kingfisher Flex robotic handling system was used for the remaining steps. 200 μg of MagReSyn® HILIC microspheres were equilibrated in 500 μl equilibration buffer (15% ACN, 100 mM ammonium acetate, pH = 4.5). Urine proteins were bound to HILIC microspheres by combining 400 μL of lysate with 400 μL of binding buffer (30% ACN, 200 mM ammonium acetate, pH = 4.5) and mixing. Microspheres were then washed twice in 95% ACN. On-sphere digestion was performed by incubating with 250 μL of digestion buffer (50 mM ammonium bicarbonate, 1 μg trypsin, 0.25 μg Lys-C) for 2h at 37°C. The digestion was stopped with 12 μL of 20% TFA (final concentration 1%) and the pH was verified to be between 1–2. Urine peptides were desalted and stored in the same way as saliva peptides.

### LC-MS proteomic plasma sample preparation

Plasma proteomic sample preparation for MS-based analysis was performed using an in-solution digestion protocol. 1 μL of plasma was mixed with 100 μL of solubilization buffer (100 mM Tris pH = 8.5, 10 mM TCEP, and 40 mM CAA) in a 96-well low-protein bind plate and boiled for 10 min at 95°C and 800 RPM in a Thermomixer. Samples were then sonicated for 10 min in a water bath sonicator. Assuming a 70 μg/μL starting concentration, half of the plasma protein lysate was digested with 1:100 trypsin and 1:500 lysC overnight at 37°C 800 RPM in a Thermomixer. Following overnight digestion (∼16 h) peptides were acidified to a final concentration of 1% TFA and desalted on a 3xlayer C18 disc plugged into a p200 tip. Desalted plasma peptides were stored in the same way as saliva peptides.

### LC-MS measurements

Peptide concentrations were measured using the Lunatic microfluidic platform at 260nm/280nm absorbance, and 200 ng was loaded onto equilibrated Evotips (Evosep). Peptides were separated on a Pepsep 8 cm, 150 μM ID column packed with C18 beads (1.5 μm) using an Evosep ONE (Evosep) HPLC system applying the default 60-SPD (60 samples per day) method. Column temperature was maintained at 35°C. Following elution, peptides were injected via a CaptiveSpray source and 20-μm emitter into a timsTOF HT MS (Bruker) operated in diaPASEF mode. MS data were collected over a 100-1700 m/z range. A modified version of the short-gradient method was used, which included 8 diaPASEF scans with three 25 Da windows per ramp, a mass range of 400.0-1000.0 Da, and a mobility range of 1.37-0.64 1/K0. The collision energy was decreased linearly from 45 eV at 1/K0 = 1.3 to 27 eV at 1/K0 = 0.85 Vs cm-2. Both accumulation time and PASEF ramp time were set to 100 ms. The total cycle time was 0.95 s.

### LC-MS data processing

Saliva, urine, and plasma LC-MS proteomics data were searched with Spectronaut v18.6 (Biognosys AG, Switzerland) in directDIA mode, using human protein FASTA (downloaded from UniProt 04.05.2024). Raw protein intensities were loaded into R (v4.4.0), underwent logarithmic transformation (log_2_), filtered for 50% missing intensity values, and normalized with medianScaling() function from the R package PhosR (v1.14.0)^50^. The total ion chromatogram (TIC) and intensity distributions were used to assess sample quality. 9/150 saliva samples, which included 1 entire participant and 4 different participants and time points, were removed due to poor sample quality (e.g., low TIC).

### Plasma Olink Explore HT proteomic profiling

#### Sample preparation and dilution

40 ul of plasma was manually transferred to 96-well plates and then robotically transferred to 384-well Sample Source Plates using the F.A.S.T.™ Liquid Handler (Formulatrix). External controls from the Olink Explore HT kit were added manually. Samples were diluted in four sequential steps using the F.A.S.T.™ instrument, following block-specific dilution factors: Blocks 1–4, 1:1; Block 5, 1:10; Block 6, 1:100; Block 7, 1:1,000; and Block 8, 1:100,000. Dilution plates were prepared with dragonfly® discovery (SPT Labtech).

#### Incubation, PCR amplification, library preparation and sequencing

Eight incubation mixes containing paired antibody-DNA conjugates were added to the diluted samples and incubated overnight at 4°C. Following overnight incubation, index primers and PCR mix were added using the F.A.S.T.™ and dragonfly® discovery instruments, respectively.

PCR was conducted on a ProFlex™ 384-well PCR System (Applied Biosystems™). PCR products were pooled into eight block-specific libraries, purified using magnetic beads, and assessed with automated electrophoresis on the TapeStation 4200 (Agilent). Equal volumes from each block were combined to create a single Olink Library per sample for sequencing. Libraries were denatured and sequenced using the Illumina NovaSeq™ 6000 platform with the S4 Reagent Kit (35 cycles) and the Olink-custom NovaSeq recipe.

#### Data processing and storage

Sequencing reads were processed with Olink® ngs2counts (v4.5.0), and quality control and NPX calculation were performed using Olink® NPX Explore HT software. Final NPX values were stored in Parquet format and used for downstream analysis in R.

### Quantification and statistical analysis Statistical analysis

Linear mixed models were used to test for changes in protein and plasma metabolic marker abundance over time relative to pre-exercise. This was accomplished by performing mean scaling, which involved computing the mean and SD for each protein at pre-exercise, then taking each protein from all samples/timepoints and subtracting the pre-exercise mean and dividing by the pre-exercise SD as previously described^30^. This was done for the Discovery cohort as well as the Replication cohort. For the Replication cohort, each exercise mode was used to scale post-exercise samples to baseline. The normalized data was used for further statistical modelling.

The R package lme4 (v1.1-35.5)^51^ was used for all mixed effect models and all statistical tests used the Benjamini–Hochberg procedure^52^ to control FDR at 5% (*q* < 0.05) for multiple comparisons. A linear mixed model was applied to examine time-dependent effects on protein abundance for the Discovery cohort. Time was used as a fixed effect, participant was used as a random effect to account for repeated measures, and the model was adjusted for covariates such as age and VO_2max_. Residuals were assessed and used to examine variability not explained by the time model. For variance decomposition analysis, pre-exercise variance (between-subject variability) and residual variance (within-subject variability across time) were also estimated using linear mixed models without adjusting for covariates. The relative contributions of these variance components were examined to identify proteins with robust exercise responses despite dynamic baseline heterogeneity.

For the Replication cohort, we again performed mean scaling relative to pre-exercise levels within each exercise mode and used mixed-effect models to examine sex or exercise mode-differential effects by testing time × sex and time x exercise mode interactions and main effects for time, sex, and exercise modes. Time was used as a fixed effect, participant ID was used as a random effect to account for repeated measures, and the model was adjusted for covariates such as age and total fat mass. Given significant main effects or interactions, we further explored individual models stratified by exercise mode or sex.

### Dimensionality reduction and visualization using t-SNE

We applied t-distributed Stochastic Neighbor Embedding (t-SNE) to body fluid proteomic data using the Rtsne package in R (v0.17) to visualize sample clustering. Only proteins quantified in 100% of samples were included. Parameters were set to 2 dimensions (dims), perplexity = 30, maximum iterations to 500, and random seed to 123. The resulting two-dimensional embeddings were assigned distinct colors for each participant, ensuring clear differentiation between the 25 participants samples.

### Clustering proteome temporal dynamics

Proteins significantly associated with time (*q* < 0.05) were extracted, and their z-scores were computed across time points based on Beta coefficients and standard errors. Clustering was performed by fitting a noise-augmented von Mises-Fisher mixture model with the navmix package (v0.2.1)^29^, enabling identification of clusters of proteins based on their temporal expression profiles. Bayesian information criterion (BIC) was used to determine the optimal cluster number for each body fluid. The clustering results were merged with the linear model output for further downstream analysis/plotting.

### Pathway and Gene Ontology enrichment analysis

Functional enrichment analysis was performed using the gost() function from gprofiler2 (v0.2.3)^53^ and enrichGO() function from clusterProfiler (v4.12.6), leveraging Kyoto Encyclepedia of Genes and Genomes (KEGG), Reactome, and GO:ALL. The background was set as the total proteome of the specific body fluid being examined. Enrichment analysis was conducted for (i) all significant proteins and (ii) each protein cluster separately and (iii) for hub proteins, exercise mode-specific proteins, and proteins with a sex by time interaction.

### Tissue and cell type enrichment analysis

Tissue and cell type-specific enrichment was performed as previously described^30^, with some modifications. Tissue and cell type specificity information were retrieved from https://www.proteinatlas.org/ (HPA, v24.0)^24^. Results from linear mixed models were integrated with HPA annotations by matching protein Uniprot identifiers. Proteins were categorized based on statistical significance (e.g., significant vs. non-significant) and temporal clustering. For each tissue and cell type, we constructed contingency tables comparing the number of proteins in each category (e.g., significant/cluster specific vs. all other proteins) that associated with a specific tissue/cell type against those not associated. Tissue or cell type specific proteins were determined based on being ‘enriched’ (mRNA expression 4 times higher than any other tissues/cell types), ‘enhanced’ (mRNA expression 4-fold higher than average of other tissues/cell types), or ‘group enriched’ (mRNA expression in 2-5 tissues/2-10 cell types is 4-fold higher than any other tissues/cell types). We applied a two-sided Fisher’s exact test to these contingency tables to assess the enrichment of sets of proteins within specific tissues or cell types.

### Disease enrichment analysis

Disease enrichment was performed similarly to the tissue/cell type enrichment. Briefly, protein incident disease associations were accessed and downloaded from https://proteome-phenome-atlas.com/ (UKBB protein-disease atlas)^27^ and split into increasing and decreasing risk based on having a hazard ratio (HR) > or < 1, respectively. Proteins from each comparison were mapped to known incident disease associations. For each unique disease, we constructed contingency tables comparing the number of proteins in each cluster or sex/exercise mode-specific proteins (e.g., cluster vs. all other proteins) that associated with a specific disease against those not associated. A two-sided Fisher’s exact test using the fisher.test() function was performed to assess the enrichment of proteins in disease categories depending on whether they were increasing (HR>1) or decreasing (HR <1) risk of incidence.

### Integrative body fluid analysis

#### Data preparation

We utilized the BioNERO (v3.21) and WGCNA (v1.72-5) packages in R for weighted protein co-expression network analysis. The dataset comprised log2-transformed protein expression values from body fluid samples for time points where all body fluids were sampled (pre-, 0 h, and 24 h post-exercise).

#### Data standardization and preparation

To ensure comparability across body fluids, we standardized the expression data by applying z-score normalization to each protein’s expression profile across samples. We addressed missing values by replacing them with zeros, utilizing the replace_na function from the BioNERO package, which assumes no difference in the mean in missing data. Next, we estimated the connectivity (Z.K standardized connectivity) to identify and remove outlier samples. Three samples with a Z.K score below 2 were considered outliers and excluded from further analysis.

#### Network construction

We constructed a weighted gene co-expression network using the WGCNA framework. The soft-thresholding power (β) was determined based on the scale-free topology criterion, aiming for a signed hybrid network type. The adjacency matrix was computed using Pearson correlation, and modules were identified through hierarchical clustering of the topological overlap matrix, followed by dynamic tree cutting.

#### Hub protein identification

Within each time point, hub proteins were identified based on their intramodular connectivity. Proteins with high connectivity were considered central to their respective networks. The top 500 hub proteins were extracted and used for ORA.

#### Module preservation analysis

We assessed the preservation of identified modules across different time points with permutation testing 500 times and compared module structures to evaluate their stability over time.

#### Correlation analysis

For each protein detected in multiple fluids, pairwise Spearman correlation analyses were conducted to assess co-expression patterns at each time point. We classified proteins based on the following criteria:

- Consistently high correlation (Both > 0.5): Proteins exhibiting positive correlations (ρ > 0.5) at both “pre” and “0 h” time points across body fluids.
- Consistently low correlation (Both < -0.5): Proteins showing negative correlations (ρ < -0.5) at both time points across body fluids.
- Divergent correlation – Top-left quadrant: Proteins with a positive correlation at “Pre” (ρ > 0.4) and a negative correlation at “0 h” (ρ < -0.4) across body fluids.
- Divergent correlation – Bottom-right quadrant: Proteins with a negative correlation at “Pre” (ρ < -0.4) and a positive correlation at “0 h” (ρ > 0.4) across body fluids.
- Shift from strong to neutral correlation: Proteins with a strong correlation across body fluids (|ρ| > 0.5) at one time point and a near-zero correlation (|ρ| < 0.1) at the other.

### Genetic analysis and graph visualization

We tested for colocalization of the proteins measured with the Olink panel and a full set of traits and diseases included on the Open Targets platform^26^. Colocalization computes the posterior probabilities (PP) of five hypotheses: the non-presence of a genetic association signal in the locus (H_0_); the presence of a genetic signal only for trait 1 (H_1_), e.g. for BMI-only; and only for trait 2 (H_2_) e.g. for the pQTL-only; the presence of a genetic signal for both traits, but with different causal variants (H_3_); and the presence of a genetic signal for both traits, which share the same causal variant (H_4_). We determined the presence of colocalization for cases where H_4_ is greater or equal to 0.8. We retrieved colocalization results from the Open Targets platform^26^ using the otargen R package^54^. Traits were filtered based on evidence of being modifiable through physical activity. Networks were visualized in graphs made with the R package igraph (v2.0.3). Approved drugs that target known exercise-regulated proteins were downloaded from https://go.drugbank.com/ (DrugBank v5.1.13)^33^ and used to make druggable sub-graphs for the most prominent trait categories.

### Resource availability

All results are made available through an interactive Shiny application https://cbmr.ku.dk/research/research-groups/deshmukh-group/shiny-apps/. Proteomics data have been deposited to the ProteomeXchange Consortium via the PRIDE^55^ partner repository with the dataset identifier PXD063954, which will be publicly available upon publication. All original code has been deposited to GitHub at https://github.com/fpm-cbmr/bodyfluid-exerome and will be publicly available on the date of publication. Any additional information required to reanalyze the data reported in this paper is available from the lead contact upon request.

## Acknowledgements

The authors would like to acknowledge Juleen R. Zierath and Steffen H. Raun for providing critical feedback on the manuscript and the Rodent Metabolic Phenotyping Platform at the Novo Nordisk Foundation Center for Basic Metabolic Research (CBMR) for their technical expertise and support in hosting our Shiny Application.

## Author contributions

Conceptualization: ASD, BKP, NZJ, and NK. Data curation: NK, NZJ, and NSN. Formal analysis: NK, JK, VD, and RAJS. Funding acquisition: ASD and BKP. Investigation: NK, JK, DS, CBL, NZJ, CD, NSN. Methodology: NK, SR, LO, ASD, and DS. Project administration: NK, ASD, RJFL, NZJ, MRL, CD, NSN. Resources: ASD, RJFL, and BKP. Software: RMJ and NK. Supervision: ASD and BKP. Visualization: NK and RMJ. Writing – original draft: NK, JK and ASD. Writing – review and editing: All authors.

## Supplementary Figures

**Figure S1.**
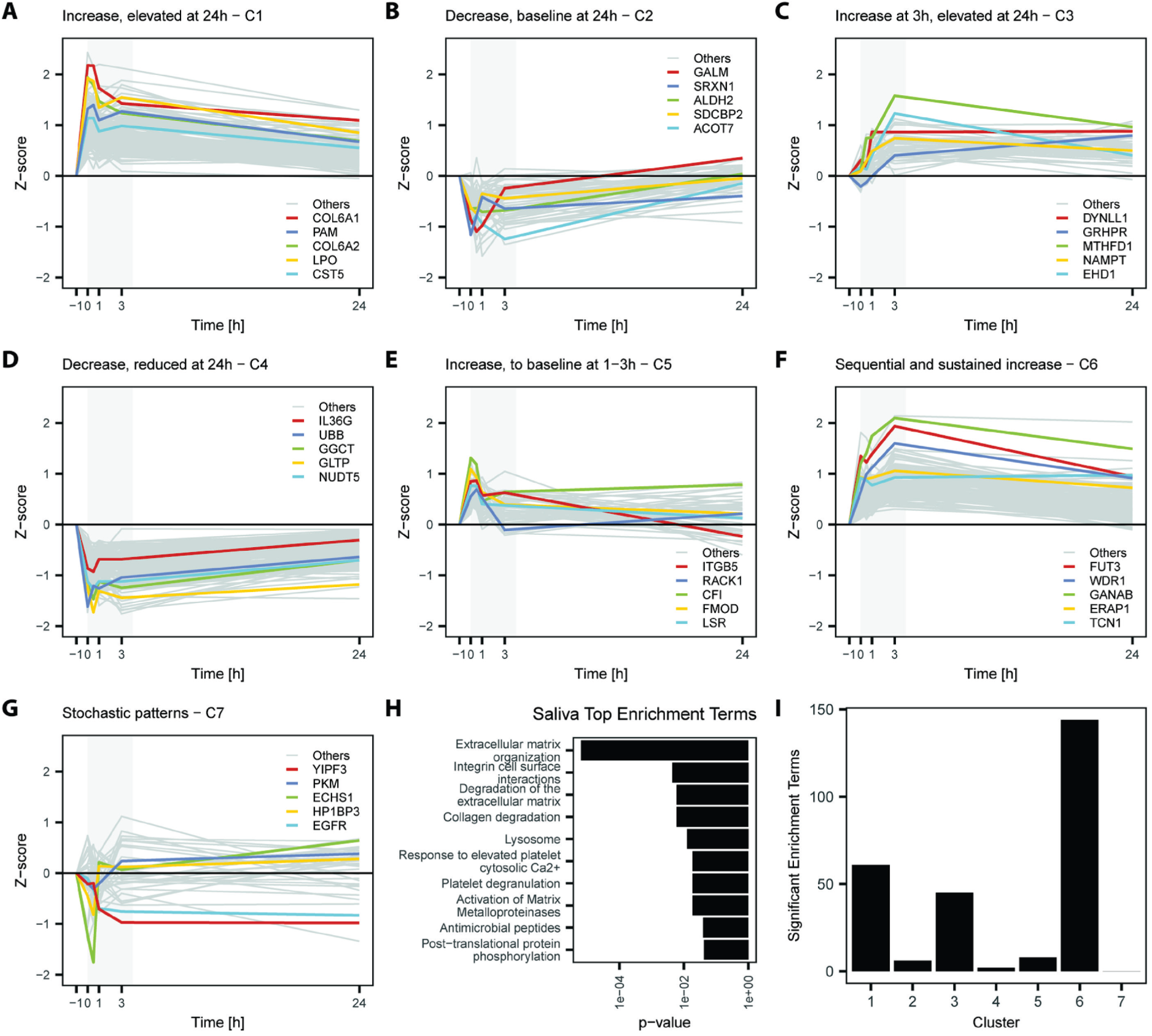
Temporally resolved exercise-induced salivary proteome dynamics. **A-G**) Temporally resolved clusters of salivary proteome dynamics with the 5 lowest p-values labeled in colors and other significant proteins in grey (associated with Figure 2D). **H)** Top 10 most significant terms from overrepresentation analysis with all 601 significantly regulated proteins. **I)** Number of significant enriched terms from cluster-specific over-representation analysis. (Associated with Figure 2).

**Figure S2.**
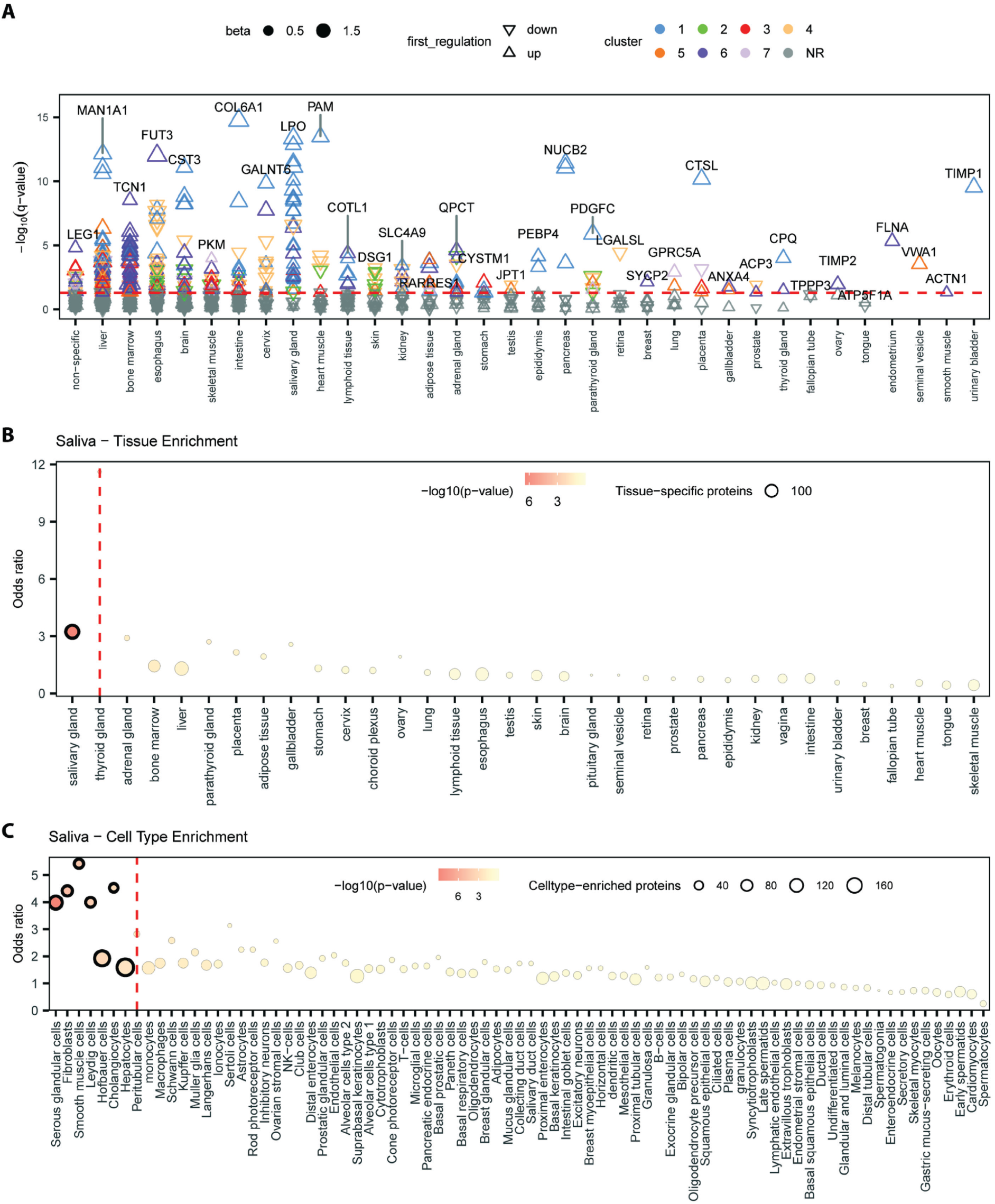
Tissue and cell type-specific contribution to the exercise-regulated salivary proteome. **A**) Salivary proteins binned into tissues based on evidence of tissue enrichment from the Human Protein Atlas (proteinatlas.org)^24^. **B)** Results from fisher exact test on tissue and **C)** cell type contribution to the 601 exercise-regulated salivary proteins. (Associated with Figures 2G and 2H).

**Figure S3.**
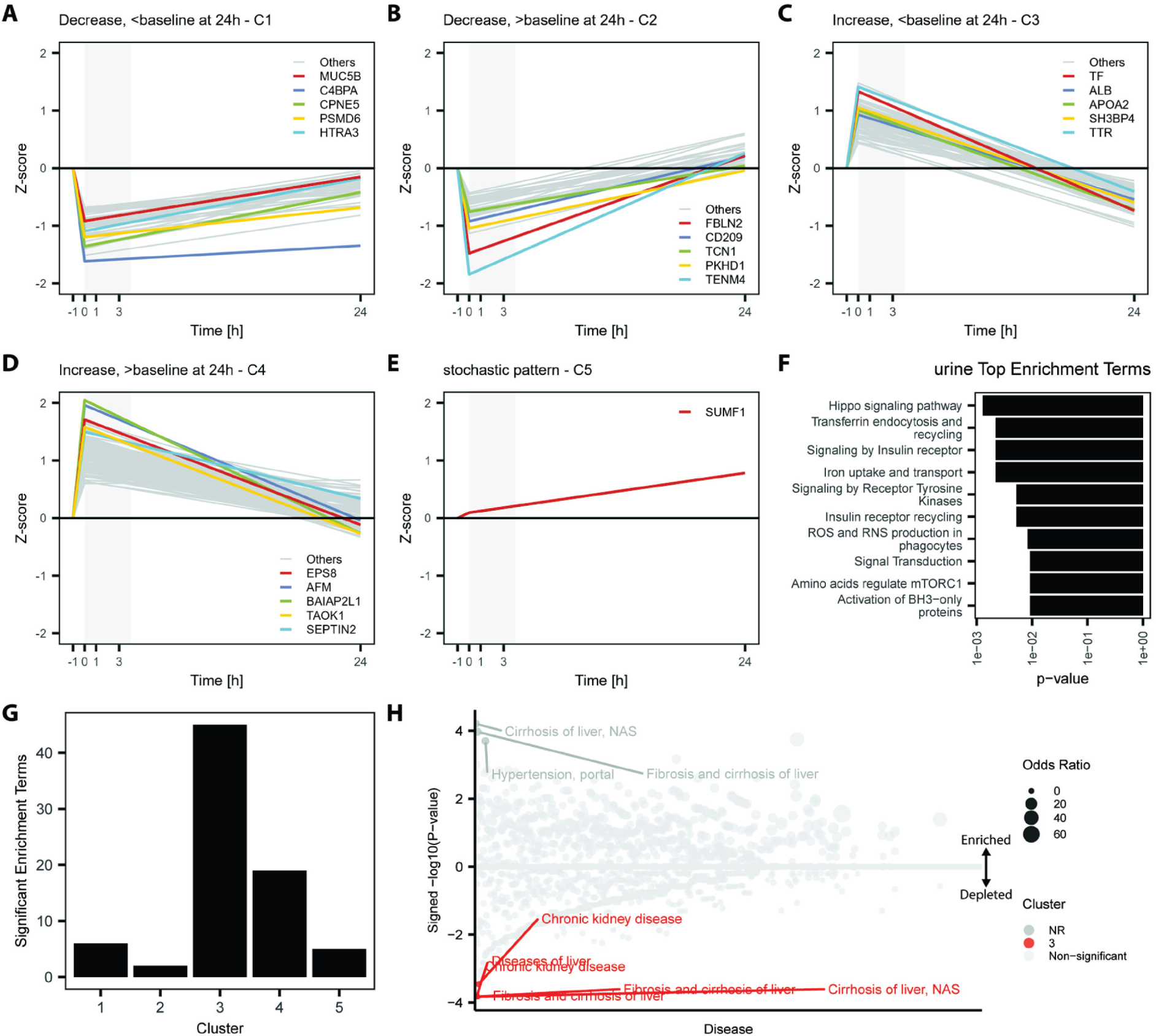
Temporally resolved exercise-induced urinary proteome dynamics. **A-E**) Temporally resolved clusters of exercise-induced urinary proteome dynamics with 5 lowest p-values labeled in colors and other significant proteins in grey. **F)** Top 10 most significant terms from over-representation analysis with all 601 significantly regulated proteins. **G)** Number of significant enriched terms from cluster-specific over-representation analysis (associated with Figure 3E). **H)** Protein incident-disease higher risk (HR>1) enrichment analysis for each cluster. (Associated with Figure 3).

**Figure S4.**
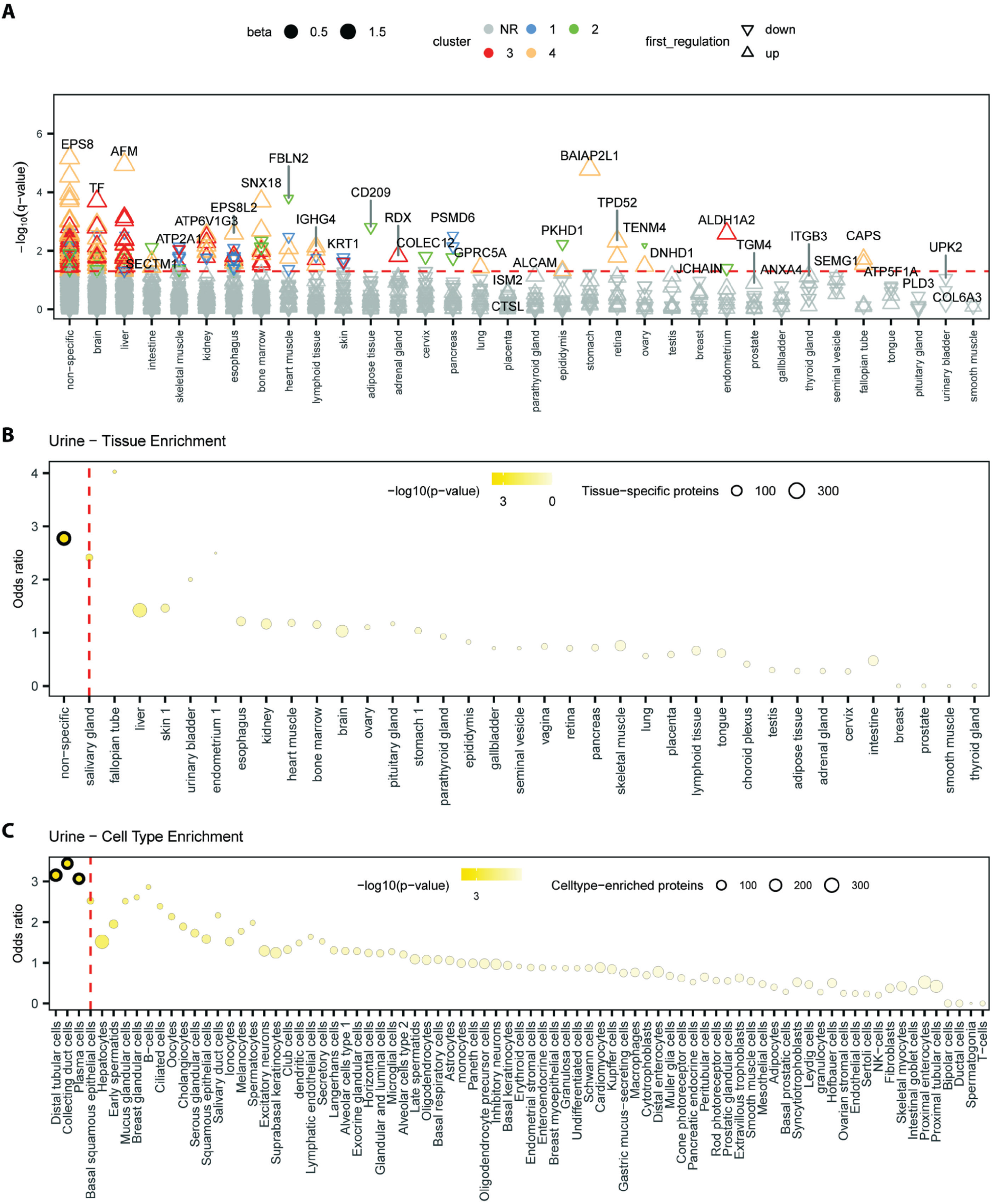
Tissue and Cell Type-Specific Contribution to the Exercise-Regulated Urinary Proteome. **A**) Urinary proteins binned into tissues based on evidence of tissue enrichment from the Human Protein Atlas (proteinatlas.org)^24^. **B)** Results from Fisher’s exact test on tissue and **C)** cell type contribution to the 272 exercise-regulated urinary proteins. (Associated with Figures 3G and 3H).

**Figure S5.**
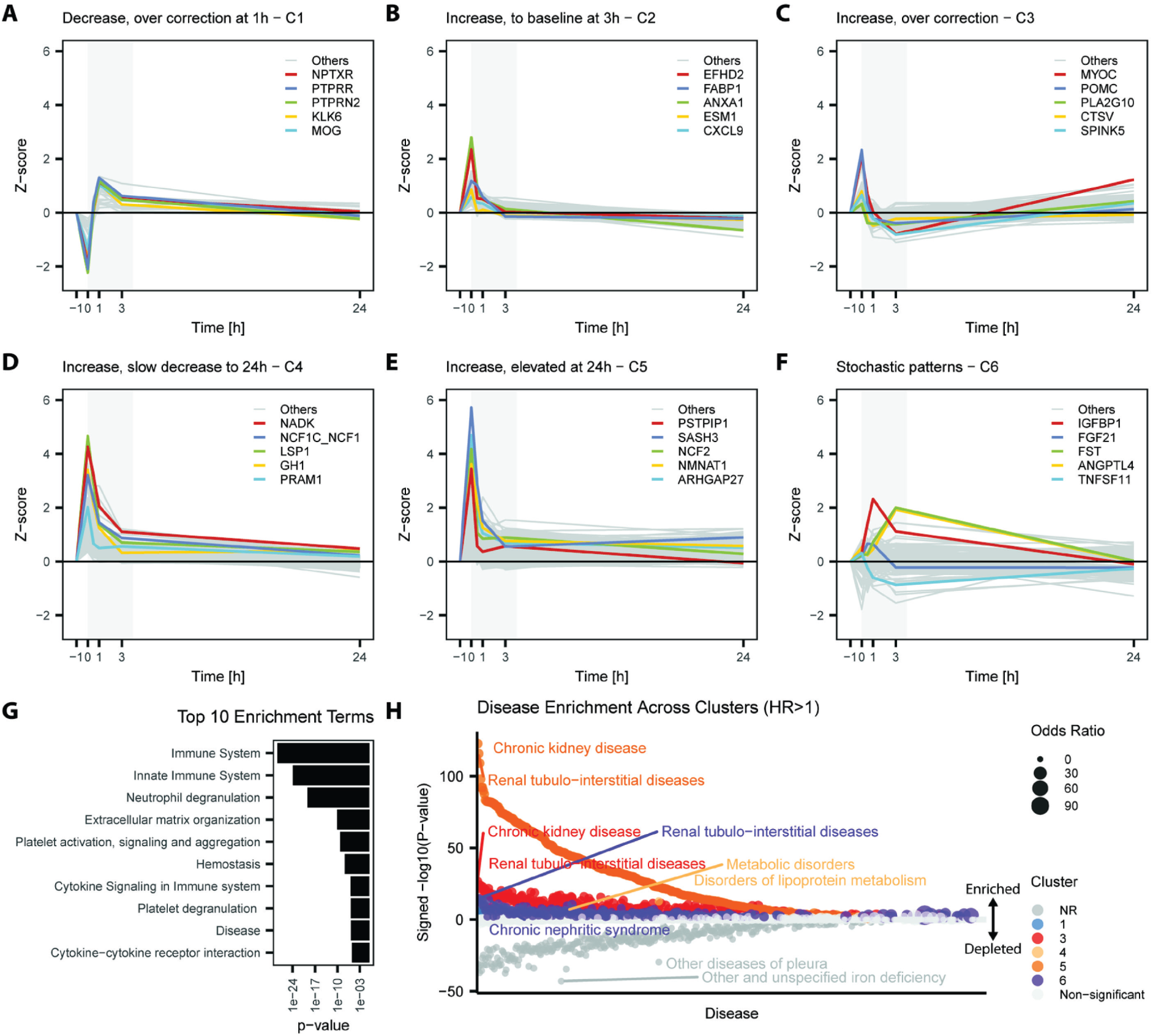
Temporally resolved exercise-induced plasma proteome dynamics. **A-F**) Temporally resolved clusters of plasma proteome dynamics with the 5 lowest p-values labeled in colors and other significant proteins in grey (q < 0.05). **G)** Top 10 most significant terms from over representation analysis with all 2156 significantly regulated proteins. **H)** Protein incident-disease higher risk (HR>1) enrichment analysis for each cluster. (Associated with Figure 4).

**Figure S6.**
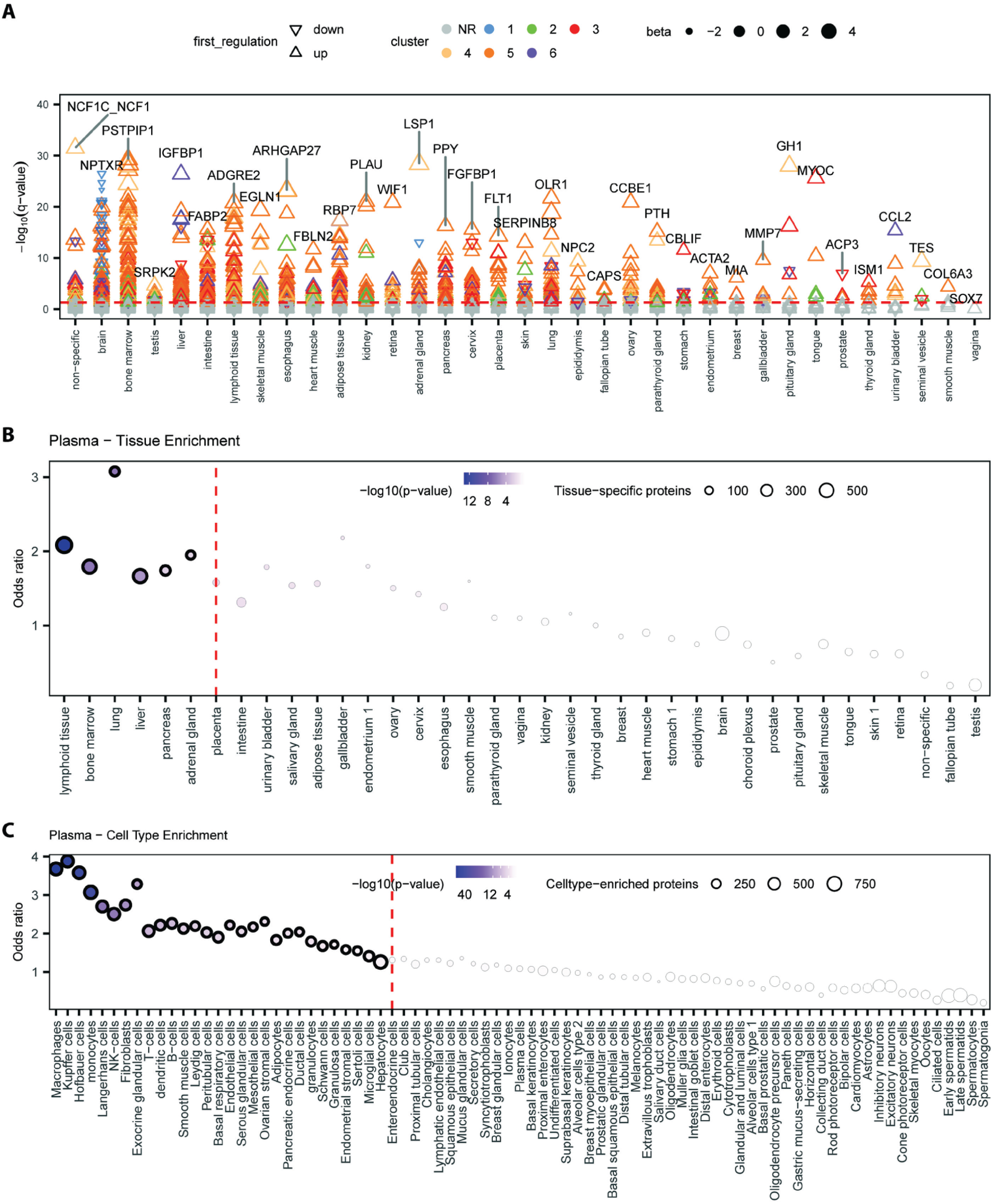
Tissue and cell type-specific contribution to the exercise-regulated plasma proteome. **A**) Plasma proteins binned into tissues based on evidence of tissue enrichment from the Human Protein Atlas (proteinatlas.org)^24^. **B)** Results from Fisher’s exact test on tissue and **C)** cell type contribution to the 2,156 exercise-regulated plasma proteins. (Associated with Figures 4G and 4H).

**Figure S7.**
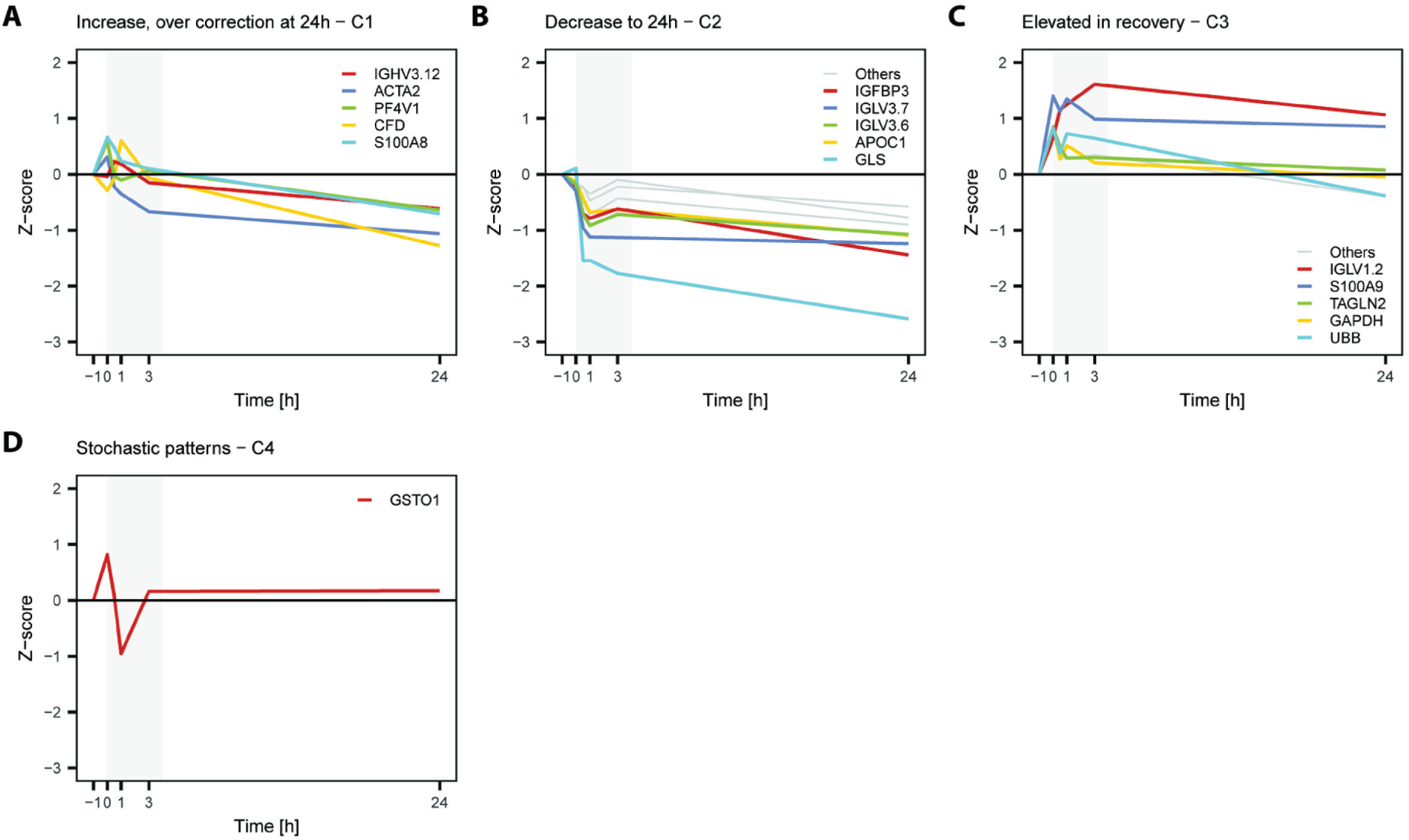
Plasma proteome response to acute exercise measured with lc-ms/ms-based proteomics. **A-D**) Temporally resolved clusters of plasma proteome dynamics with the 5 lowest p-values labeled in colors and other significant proteins in grey (q < 0.05) from identified clusters.

**Figure S8.**
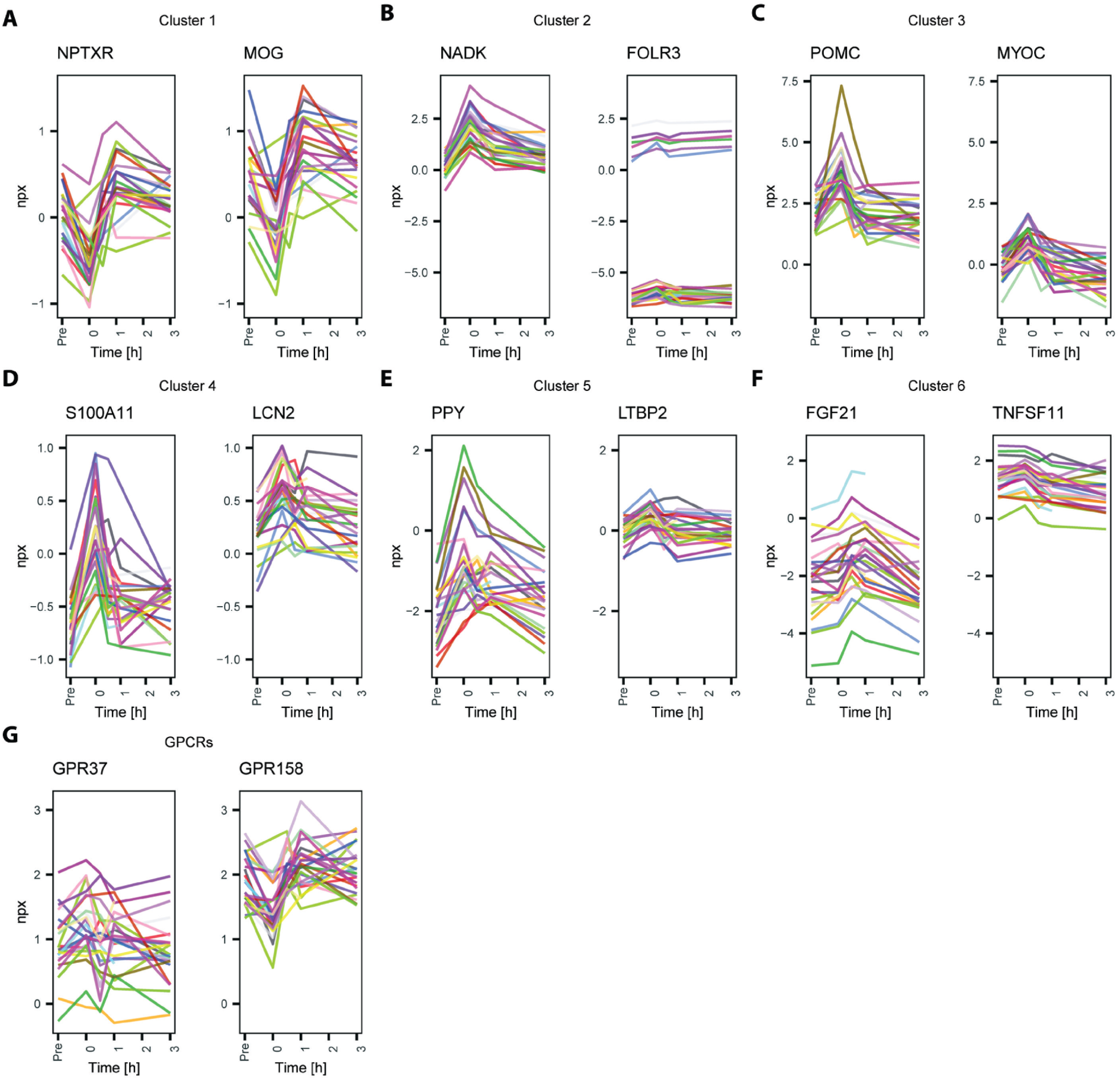
Examples of specific proteins with robust responses across participants for each plasma proteome dynamic cluster. **A-F**) NPX values plotted for each participant for two protein examples from each cluster (1-6) and **F)** two GPCRs that were regulated by acute exercise.

## Supplementary Table legends

**Table S1. Cohort characteristics**.

**A)** Discovery and Replication cohorts’ inclusion and exclusion criteria (Associated with Figures 1A and 6A). **B)** Discovery and Replication cohorts’ characteristics (Associated with Figures 1A and 6A). **C)** Discovery cohort baseline blood biochemistry (Associated with Figures 1A-1C).

**Table S2. Linear mixed model results**.

**A)** Discovery cohort exercise induced metabolic markers and cytokine linear mixed model results (Associated with Figure 1). **B)** Discovery cohort salivary results from linear mixed models for time effects (Associated with Figures 2, S1, and S2). **C)** Discovery cohort urinary results from linear mixed models for time effects (Associated with Figures 3, S3, and S4). **D)** Discovery cohort plasma Olink results from linear mixed models for time effects (Associated with Figures 4 and S5-S9). **E)** Discovery cohort plasma LC MS-based results from linear mixed models for time effects (Associated with Figure S7). **F)** Replication cohort plasma Olink results from linear mixed models for time and exercise mode (group) effects and their interaction (Associated with Figure 6B). **G)** Replication cohort plasma Olink combined individual exercise mode results from linear mixed models for time effects (Associated with Figure 6D). **H)** Replication cohort plasma Olink results from linear mixed models for sex, exercise mode, and time effects and their interactions (Associated with Figure 6F).

**Table S3. Overrepresentation analysis results**.

**A)** Discovery cohort salivary all significant proteins overrepresentation analysis results (Associated with Figure S1H). **B)** Salivary cluster-specific overrepresentation analysis results (Associated with Figure 2E). **C)** Discovery cohort urinary all significant proteins overrepresentation analysis results (Associated with Figure S3). **D)** Discovery cohort urinary cluster-specific overrepresentation analysis results (Associated with Figure 3E). **E)** Discovery cohort plasma Olink all significant proteins overrepresentation analysis results (Associated with Figure S5). **F)** Discovery cohort plasma Olink cluster-specific overrepresentation analysis results (Associated with Figure 4E). **G)** Overrepresentation analysis of WGCNA unique top hub proteins per time point (Associated with Figure 5E). **H)** Replication cohort plasma Olink cluster overrepresentation analysis results (Associated with Figure 6E). **I)** Replication cohort plasma Olink significant sex by time proteins overrepresentation analysis results (Associated with Figure 6G).

**Table S4. Disease enrichment results**.

**A)** Increased incident disease risk enrichment analysis for salivary proteome exercise-induced dynamic clusters (Associated with Figure 2J). **B)** Decreased incident disease risk enrichment analysis for salivary proteome exercise-induced dynamic clusters (Associated with Figure 2J).

**C)** Increased incident disease risk enrichment analysis for urinary proteome exercise-induced dynamic clusters (Associated with Figure S3H). **D)** Decreased incident disease risk enrichment analysis for urinary proteome exercise-induced dynamic clusters (Associated with Figure S3H). **E)** Increased incident disease risk enrichment analysis for plasma proteome exercise-induced dynamic clusters (Associated with Figure S5H). **F)** Decreased incident disease risk enrichment analysis for plasma proteome exercise-induced dynamic clusters (Associated with Figure 4I) **G)** Increased incident disease risk enrichment analysis for sex by time and exercise mode specific proteins (Associated with Figure 6H). **H)** Decreased incident disease risk enrichment analysis for sex by time and exercise mode specific proteins (Associated with Figure 6H).

**Table S5. Colocalization analysis results for cis s/e/pQTLs and traits available in open targets** (Associated with Figure 7).

